# Wnt Regulation: Exploring Axin-Disheveled interactions and defining mechanisms by which the SCF E3 ubiquitin ligase is recruited to the destruction complex

**DOI:** 10.1101/847384

**Authors:** Kristina N. Schaefer, Mira Pronobis, Clara E. Williams, Shiping Zhang, Lauren Bauer, Dennis Goldfarb, Feng Yan, M. Ben Major, Mark Peifer

**Affiliations:** Curriculum in Genetics and Molecular Biology, University of North Carolina at Chapel Hill, Chapel Hill, NC 27599, USA; Department of Biology, University of North Carolina at Chapel Hill, CB#3280, Chapel Hill, NC 27599-3280, USA; Department of Cell Biology and Physiology, University of North Carolina at Chapel Hill, Chapel Hill, NC 27599, USA; Lineberger Comprehensive Cancer Center, University of North Carolina at Chapel Hill, Chapel Hill, NC 27599, USA; Department of Cell Biology and Physiology, Washington University School of Medicine, St. Louis, MO 63110, USA; Institute for Informatics, Washington University School of Medicine, St. Louis, MO 63110, USA; Department of Otolaryngology, Washington University School of Medicine, St. Louis, MO 63110, USA

## Abstract

Wnt signaling plays key roles in embryonic development and adult stem cell homeostasis, and is altered in human cancer. Signaling is turned on and off by regulating stability of the effector β-catenin. The multiprotein destruction complex binds and phosphorylates β-catenin, and transfers it to the SCF-TrCP E3-ubiquitin ligase, for ubiquitination and destruction. Wnt signals act though Dishevelled to turn down the destruction complex, stabilizing β-catenin. Recent work clarified underlying mechanisms, but important questions remain. We explore β-catenin transfer from the destruction complex to the E3 ligase, and test models suggesting Dishevelled and APC2 compete for association with Axin. We find that Slimb/TrCP is a dynamic component of the destruction complex biomolecular condensate, while other E3 proteins are not. Recruitment requires Axin and not APC, and Axin’s RGS domain plays an important role. We find that elevating Dishevelled levels in *Drosophila* embryos has paradoxical effects, promoting the ability of limiting levels of Axin to turn off Wnt signaling. When we elevate Dishevelled levels, it forms its own cytoplasmic puncta, but these do not recruit Axin. SIM imaging in mammalian cells suggests that this may result by promoting Dishevelled: Dishevelled interactions at the expense of Dishevelled:Axin interactions when Dishevelled levels are high.

## Introduction

During embryonic development cells must choose fate based on their position within the unfolding body plan. One key is cell-cell signaling, by which cells communicate positional information to neighbors and ultimately direct downstream transcriptional programs. A small number of conserved signaling pathways play an inordinately important role in these events in all animals. These include the Hedgehog, Notch, Receptor Tyrosine kinase, BMP/TGFβ, and Wnt pathways, which influence development of most tissues and organs (Basson, 2012). These same signaling pathways regulate tissue stem cells during tissue homeostasis, and play critical roles in most solid tumors. Due to their powerful effects on cell fate and behavior, evolution has shaped dedicated machinery that keeps each signaling pathway definitively off in the absence of ligand.

In the Wnt pathway, signaling is turned on and off by regulating stability of the key effector β-catenin (βcat; reviewed in (Nusse and Clevers, 2017). In the absence of Wnt ligands, newly synthesized βcat is rapidly captured by the multiprotein destruction complex. Within this complex, the protein Axin acts as a scaffold, recruiting multiple partners. Axin and Adenomatous polyposis coli (APC) bind βcat, and present it to the kinases Casein kinase 1 (CK1) and Glycogen synthase kinase 3 (GSK3) for sequential phosphorylation of a series of N-terminal serine and threonine residues on βcat.

It has become increasingly clear that the destruction complex is not a simple four-protein entity. Instead, Axin directs assembly of destruction complex proteins into what the field originally described as “puncta”. We now recognize these as examples of supermolecular, non-membrane bound cellular compartments (reviewed in (Gammons and Bienz, 2017; Schaefer and Peifer, 2019), referred to as biomolecular condensates (Banani *et al*., 2017). Condensate formation is driven by Axin polymerization via its DIX domain, by APC function, and by other multivalent interactions e.g. (Fagotto *et al*., 1999; Kishida *et al*., 1999; Cliffe *et al*., 2003; Schwarz-Romond *et al*., 2007a; Faux *et al*., 2008; Mendoza-Topaz *et al*., 2011; Kunttas-Tatli *et al*., 2014; Thorvaldsen *et al*., 2015) (Pronobis *et al*., 2015; Pronobis *et al*., 2017).

Once the destruction complex templates βcat phosphorylation, the most N-terminal phosphorylated serine forms part of the core of a recognition motif for an F-box protein, mammalian βTrCP/Drosophila Slimb, which is part of a Skp-Cullin-F-box (SCF)-class E3 ubiquitin ligase. This E3-ligase ubiquitinates βcat for proteasomal destruction (Stamos and Weis, 2013). While down-regulation of βcat levels via protein degradation is a key function of the destruction complex our understanding of *how* βcat is transferred from the complex to the SCF E3 ligase is a key unanswered question.

Two classes of models seem plausible. In the first class of models, the E3 ligase is a physical entity separate from the destruction complex –this would fit with the many roles for the SCF^Slimb^ E3 ligase, which binds and ubiquitinates diverse phospho-proteins, ranging from the Hedgehog effector Ci/Gli to the centrosome assembly regulator PLK4 (Robertson *et al*., 2018). However, given the abundance of cellular phosphatases, this model has a potential major problem. Phosphorylated βcat released free from the destruction complex into the cytoplasm would likely be rapidly dephosphorylated, preventing its recognition by the E3 ligase. Consistent with this, earlier work revealed that APC helps prevent βcat dephosphorylation within the destruction complex (Su *et al*., 2008). In a second class of models, the SCF^Slimb^ E3 ligase might directly dock on or even become part of the destruction complex, either by direct interaction with destruction complex proteins or using phosphorylated βcat as a bridge. In this model, once βcat is phosphorylated it could be directly transferred to the E3 ligase, thus preventing dephosphorylation of βcat by cellular phosphatases during transit. Immuno-precipitation (IP) experiments in animals and cell culture revealed that βTrCP can coIP with Axin, APC, βcat, and GSK3, and that Wnt signals reduce Axin: βTrCP coIP (Hart *et al*., 1999; Kitagawa *et al*., 1999; Liu *et al*., 1999; Li *et al*., 2012). However, these studies did not examine whether βTrCP or other components of the E3 are recruited to the destruction complex, leaving both models an option, especially if βTrCP acts as a shuttling protein between complexes. Here we address this issue.

A second set of outstanding questions concern the mechanisms by which Wnt signaling down-regulates βcat destruction. Wnt signaling is initiated when Wnt ligands interact with complex multi-protein receptors, comprised of Frizzled family members plus LRP5/6 (reviewed in (DeBruine *et al*., 2017; Nusse and Clevers, 2017). This receptor complex recruits the destruction complex to the plasma membrane, via interaction of Axin with the phosphorylated LRP5/6 tail and with the Wnt effector Disheveled (Dvl in mammals/Dsh in Drosophila). This leads to downregulation of the destruction complex, reducing the rate of βcat destruction. Current data suggest destruction complex downregulation occurs via multiple mechanisms (reviewed in (MacDonald and He, 2012; Nusse and Clevers, 2017), some rapid and others initiated more slowly. These include direct inhibition of GSK3 by the phosphorylated LRP5/6 tail, inhibition of Axin homo-polymerization by competition with hetero-polymerization with Dsh, competition between Dsh and APC2 for access to Axin, targeting Axin for proteolytic destruction, and blockade of βcat transfer to the E3-ligase. In our recent work, we explored the role of Dsh. We found that overall protein levels of Axin, APC2 and Dsh in Drosophila embryos experiencing active Wnt signaling are within a few-fold of one another, suggesting that competition is a plausible mechanism for destruction complex down-regulation (Schaefer *et al*., 2018). The competition model is also consistent with the effects of elevating Axin levels, which makes the destruction complex more resistant to turn-down (Cliffe *et al*., 2003) (Wang *et al*., 2016; Schaefer *et al*., 2018). However, somewhat surprisingly, elevating Dsh levels had only modest consequences on cell fate choices, and Dsh only assembled into Axin puncta in cells receiving Wingless signals (Wg-Drosophila Wnt homolog; (Cliffe *et al*., 2003; Schaefer *et al*., 2018), suggesting that Dsh may need to be “activated” by Wnt signals in order to effectively compete with APC for Axin and thus mediate destruction complex downregulation. Candidate phosphorylation sites and kinases potentially involved in this activation have been identified (e.g. (Bernatik *et al*., 2011; Gonzalez-Sancho *et al*., 2013; Bernatik *et al*., 2014). Intriguingly, when Axin, APC, and Dvl were expressed in mammalian cells, potential competition between APC and Dvl for interaction with Axin was revealed (Schwarz-Romond *et al*., 2005; Mendoza-Topaz *et al*., 2011). Here we examined in vivo the effects of simultaneously altering levels of Dsh and Axin, testing aspects of the competition model, and combined this with analysis of how Dsh and Axin affect one another’s assembly into puncta in a simple cell culture model, using structured illumination super-resolution microscopy (SIM).

## Results

### A system to examine whether the destruction complex and the SCF^Slimb^ E3 ligase co-localize

The transfer of phosphorylated βcat from the destruction complex to the E3 ubiquitin ligase to begin βcat degradation is a crucial step in Wnt signaling regulation which remains incompletely understood. One key question in the field involves the mechanism by which βcat is transferred from one complex to the other. Do they each form separate structures within the cytoplasm of the cells, thus relying on either diffusion or some form of protein shuttle to move βcat between them? Or is there a “factory” for βcat destruction, containing the machinery to first phosphorylate βcat, and then to directly pass it down the assembly line to the E3 ligase? Previous work is more consistent with the latter model, as Axin and APC can co-immunoprecipitate (coIP) with mammalian βTrCP (Hart *et al*., 1999; Kitagawa *et al*., 1999; Liu *et al*., 1999; Li *et al*., 2012) and one role of APC is to protect βcat from dephosphorylation before it is ubiquitinated (Su *et al*., 2008).

We took two different approaches. First, we examined recruitment of E3 ligase components using mass spectroscopy. We used affinity-purification mass spectrometry (AP/MS) to pull down Flag-tagged APC2 that had been stably expressed in HEK293T cells. Mass spec analysis identified the known core components of the destruction complex, including βcat, alpha-catenin, GSK3, CK1, Axin1, and Axin2, as well as other proteins previously identified to interact with the destruction complex by mass spectroscopy;(.g., WTX/AMER1, plakoglobin, USP7, and CTBP; (Major *et al*., 2007; Hilger and Mann, 2012; Li *et al*., 2012)). Our list also included a number of components of different E3 ligases, including the three core components of the SCF^Slimb^ E3 ligase (Known destruction complex proteins and components of E3 ligases are summarized in Table 1 with full data set in Supplemental Table 2). Using the label-free quantification (LFQ) intensity as a measure for quantification, our data was consistent with the following hierarchical order: FBXW11=βTrCP2 > SKP1 > Cul1. However, the recovery of these proteins was less robust than that of the core destruction complex proteins. βTrCP2 and Skp1 were also identified in previous mass spectroscopy analysis with Axin as a bait, and Skp1 was also identified in APC pulldowns (Hilger and Mann, 2012). These data are consistent with the possibility that the SCF^Slimb^ E3 ligase is recruited, at least transiently, to the destruction complex.

**Table 1:** Proteins identified using affinity-purification mass spectrometry

|  | Protein | Log<sub>2</sub>(Intensity) | Unique Peptides | MS/MS Counts |
| --- | --- | --- | --- | --- |
| Destruction Complex | APC | 31.67 | 122 | 233 |
|  | CTNNB1 | 30.96 | 30 | 69 |
|  | CTNNA1 | 30.24 | 44 | 57 |
|  | JUP | 27.29 | 21 | 25 |
|  | AMER1 | 27.10 | 21 | 25 |
|  | AXIN1 | 26.33 | 16 | 21 |
|  | GSK3B | 25.39 | 8 | 11 |
|  | AXIN2 | 24.11 | 7 | 9 |
|  | CSNK1D/E | 21.66 | 2 | 1 |
| E3-ligase Components | TRIM21 | 27.96 | 17 | 24 |
|  | TRIM28 | 24.41 | 7 | 7 |
|  | FBXW11 | 23.27 | 2 | 4 |
|  | SKP1 | 22.65 | 2 | 2 |
|  | CUL1 | 21.86 | 1 | 1 |
|  | MYCBP2 | 21.53 | 5 | 5 |
|  | CUL3 | 19.49 | 1 | 1 |
|  | CDC27 | — | 1 | 1 |

To further address this issue, we transfected components of the destruction complex and of the E3 ligase into SW480 colorectal cancer cells, which are mutant for human *APC,* to determine whether they co-localize, thus suggesting recruitment of the E3 ligase to the destruction complex. For our studies we utilized the *Drosophila* proteins, which can rescue βcat destruction in this colorectal cell line (Roberts *et al*., 2011; Pronobis *et al*., 2015); Fig. 1A-C). *Drosophila* APC2 is also half the size of human APC1 and therefore easier to transfect and express in cells. To visualize the destruction complex, we tagged *Drosophila* Axin or APC2 with GFP, RFP, or Flag epitope tags (Roberts *et al*., 2011; Pronobis *et al*., 2015). When GFP-tagged APC2 (GFP:APC2) is transfected in cells alone, APC2 is found throughout the cytoplasm (Roberts *et al*., 2011); Fig 1A). In contrast, when RFP-tagged Axin is transfected alone (Axin:RFP), it forms cytoplasmic puncta, due to Axin’s ability to self-polymerize via its DIX domain (Fig 1B; (Kishida *et al*., 1999). Finally, when GFP:APC2 is expressed along with Axin:RFP, GFP:APC2 is recruited into Axin puncta (Roberts *et al*., 2011); Fig 1C). Previous studies revealed that this APC2-Axin interaction leads to larger, stabilized destruction complexes (Kunttas-Tatli *et al*., 2014; Pronobis *et al*., 2015).

**Figure 1:**
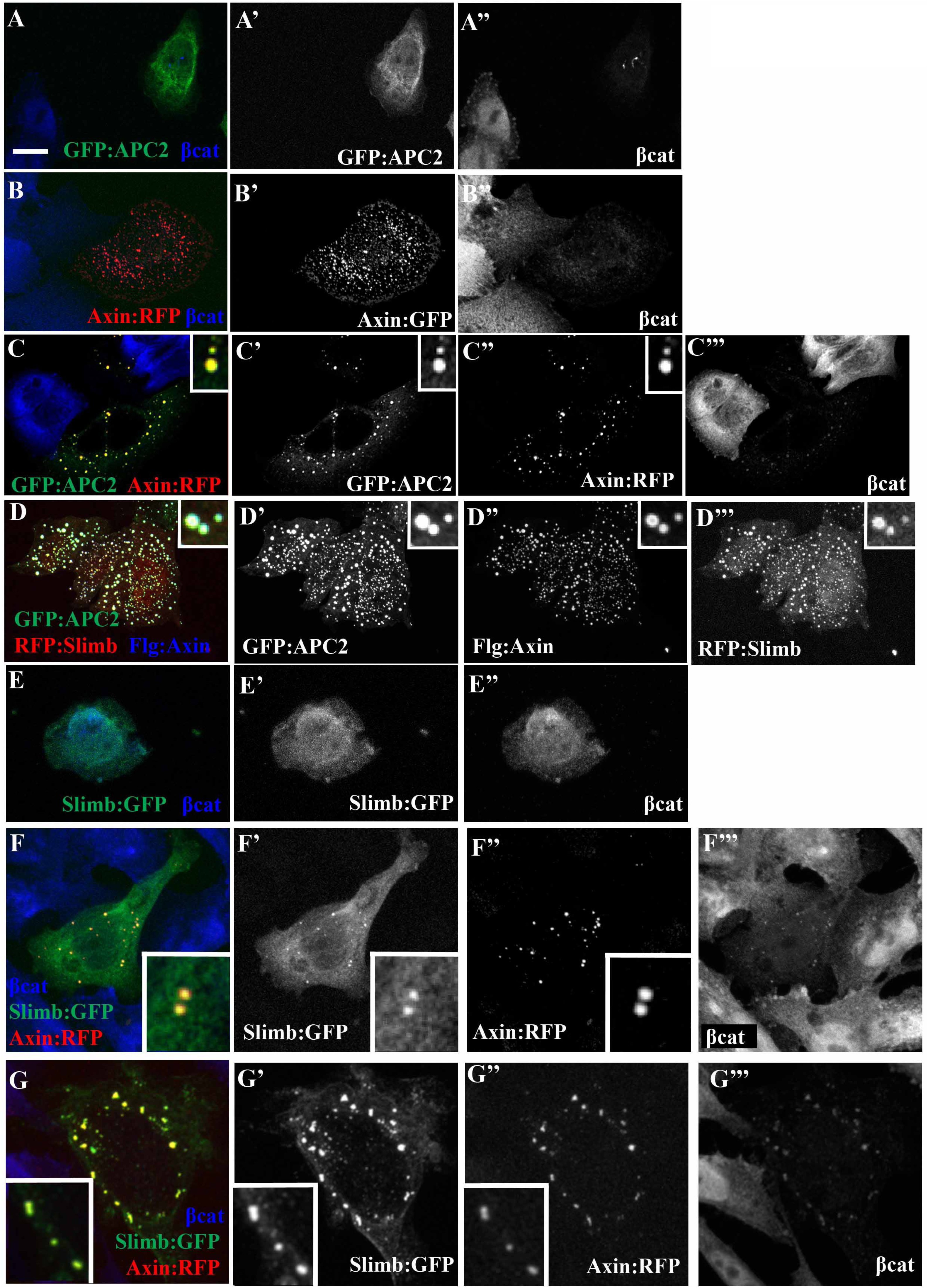
Slimb is recruited into the destruction complex by Axin. SW480 cells transfected with the indicated constructs, encoding the *Drosophila* proteins. (A) Expression of GFP:APC2 is diffuse throughout cytoplasm and nucleus. SW480 cells lack a functional human APC and thus have high levels of βcat. Addition of *Drosophila* APC2 rescues βcat destruction. (B) Axin:RFP expressed alone forms cytoplasmic puncta, due to Axin’s self-polymerization domain. (C) When co-expressed, Axin:RFP recruits GFP:APC2 into Axin puncta. (D) When co-expressed, GFP:APC2 and Flg:Axin can robustly recruit RFP:Slimb into puncta. (E) Expression of Slimb:GFP only results in diffuse localization of Slimb in the cytoplasm and nucleus. (F-G) Axin:RFP can recruit Slimb:GFP into puncta. Axin:RFP either recruits a fraction of Slimb into puncta, leaving a large cytoplasmic pool of Slimb:GFP (F) or robustly recruits most of Slimb:GFP into puncta (G). Scale bar = 10µm.

We next assessed whether Slimb has any specific localization pattern on its own in SW480 cells. We tagged *Drosophila* Slimb with either GFP, RFP or Flag tags. When Slimb was expressed alone, it was diffusely localized in both the cytoplasm and nucleus, without obvious enrichment in any subcellular structure (Fig 1E). In a few cells, there was slight enrichment of proteins in puncta near the nucleus, which may be due to the E3’s known role in regulating centrosome duplication (Wojcik *et al*., 2000). This system thus provided a platform to examine whether different components of the SCF^Slimb^ E3 ligase are recruited to the destruction complex

### Axin can recruit Slimb into the destruction complex while APC2 does not

Previous studies revealed that Axin and APC can coIP with βTrCP (Hart *et al*., 1999; Kitagawa *et al*., 1999; Liu *et al*., 1999; Li *et al*., 2012), and βTrcP2 was identified in complex with Axin by mass spectroscopy (Hilger and Mann, 2012). We first examined whether this interaction was sufficient to recruit the βTrCP homolog Slimb into destruction complex puncta. When we co-expressed both Flag:Axin and GFP:APC2 with an RFP-tagged Slimb, Slimb was robustly recruited to Axin/APC puncta (Fig 1D). However, this did not discriminate whether Axin or APC recruited Slimb.

When APC was first discovered, it was believed to be the scaffold of the destruction complex, as it binds βcat and coIPs with the kinase GSK3 (Polakis, 1997). However, subsequent work revealed that Axin is the actual destruction complex scaffold, mediating complex assembly by directly binding all core destruction complex components: APC, GSK3, CK1, and βcat (Spink *et al*., 2000; Liu *et al*., 2002; Dajani *et al*., 2003). To define whether APC and/or Axin can recruit Slimb, we co-expressed Axin plus Slimb or APC2 plus Slimb. The ability of Axin to form puncta made examining Slimb recruitment straightforward. Co-expression of RFP-tagged Axin (Axin:RFP) with GFP-tagged Slimb (Slimb:GFP) revealed that Axin robustly recruits Slimb:GFP into cytoplasmic puncta (Fig 1F-G and close-up insets; 135/140 cells examined).The degree of Slimb:GFP recruitment into Axin:RFP puncta varied from Slimb enrichment in puncta with a remaining cytoplasmic pool (Fig 1F) to nearly complete recruitment into puncta (Fig 1G).

Since APC2 has no specific localization pattern when expressed on its own (Fig 1A), it is difficult to assess whether APC2 can recruit other proteins. We therefore utilized an APC2 construct containing a mitochondrial localization signal (mito:APC2;(Roberts *et al*., 2012). Mito:APC2 is readily recruited to the mitochondria, and remains functional, as evidenced by reduction of βcat levels in SW480 cells and the ability to rescue *Drosophila* APC2 mutants (Fig 2A; (Roberts *et al*., 2012). Mito:APC2 effectively recruits Axin (Fig 2B; (Roberts *et al*., 2012). We therefore expressed GFP-tagged mito:APC2 with RFP-tagged Slimb to test whether APC2 can recruit Slimb. Mito:APC2 was unable to detectably recruit Slimb (Fig 2C; 0/10 cells examined), consistent with the idea APC2 does not directly interact with this E3 ligase component. This is consistent with previous work demonstrating that coIP of βTrCP2 and APC required co-expression of Axin (Kitagawa *et al*., 1999).

**Figure 2:**
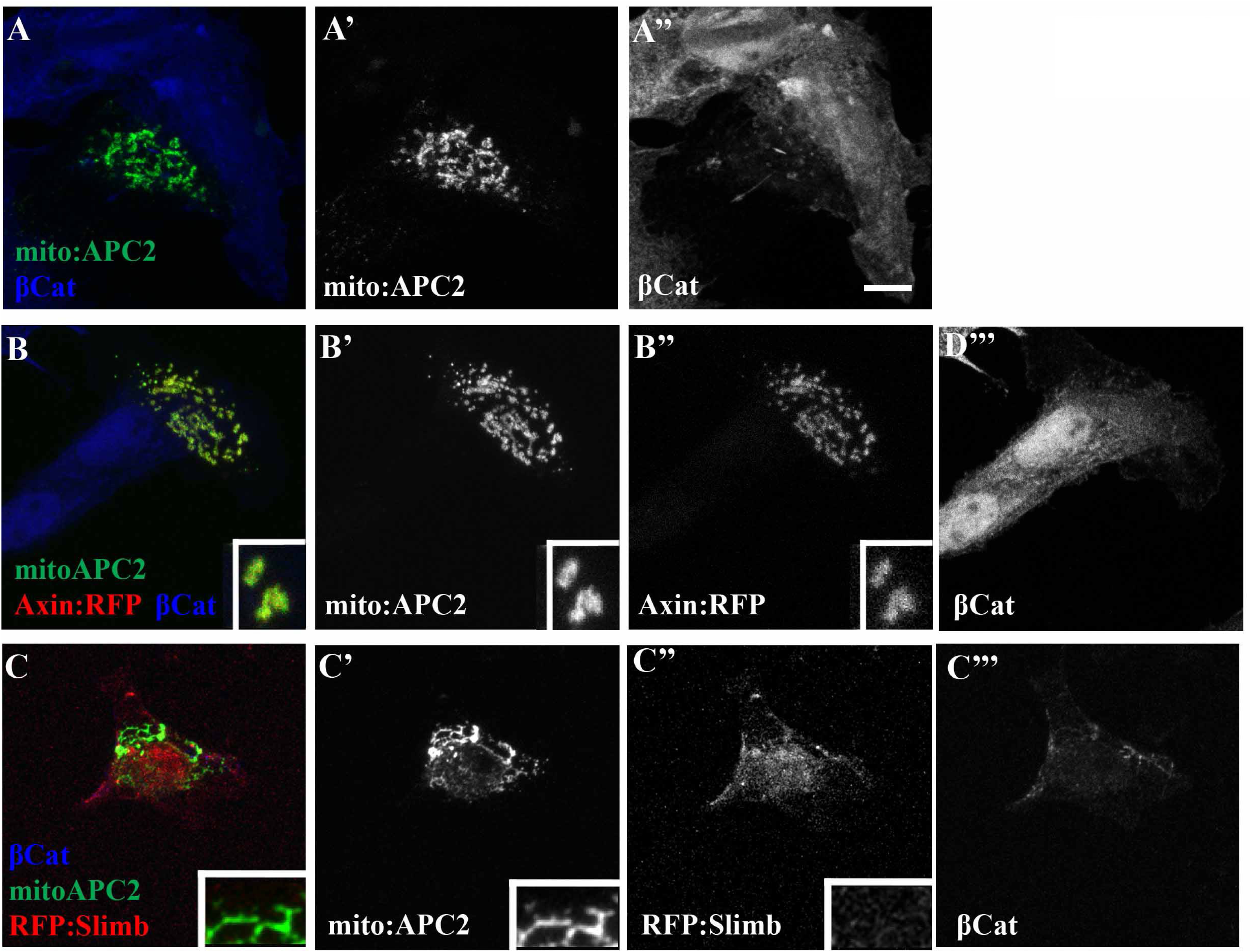
APC2 is unable to recruit Slimb. (A-C) SW480 cells transfected with the indicated constructs. (A) Expression of a GFP-tagged APC2 with a mitochondrial localization signal. Even though APC2 is immobilized at the mitochondrial membrane, it is still able to enhance βcat destruction. (B) Mito:APC2 was co-expressed with Axin:RFP. Mito:APC2 was still able to recruit Axin:RFP. (C) Mito-APC2 was expressed with RFP tagged Slimb. RFP:Slimb was not recruited by APC2. Scale bar = 10μm.

### Recruitment of other SCF^Slimb^ E3 ligase components by Axin is substantially less robust

We next explored whether the other SCF^Slimb^ E3 ligase proteins, SkpA or Cul1, are recruited into the destruction complex. To test this, we co-expressed Axin:RFP with either GFP-tagged Cul1 (GFP:Cul1) or GFP-tagged SkpA (GFP:SkpA). When expressed alone, both SkpA and Cul1 were found throughout the cytoplasm and nucleus (Fig 3A,B). We were surprised to find that while Slimb was robustly recruited to Axin puncta (Fig 1F,G), Cul1 and SkpA were not. For SkpA there was no recruitment in 48/52 cells examined (Fig 3C), and for Cul1 we observed no recruitment in 36/42 cells examined (Fig 3D). When co-localization was observed for SkpA or Cul1, it was minimal (Fig 3C inset). Consistent with previous work with the mammalian homologs, we could co-IP Axin with Slimb (Fig. 3E, middle lane; 3F, lane 2), but did not detect robust co-IP of APC2 with Slimb (Fig 3E, left lane). Similarly, we did not detect co-IP of Axin with either Cul1 or SkpA (Fig 3F, left 2 lanes). These data suggest that Axin can recruit Slimb to the destruction complex but is unable to strongly recruit the other components of the E3. We also examined whether Slimb recruitment into Axin puncta stimulated recruitment of other SCF^Slimb^ E3 ligase proteins. To do so, we co-expressed Axin, Slimb and SkpA—in this case we saw modest co-recruitment of Slimb and SkpA to Axin puncta in small subset of cases (3/20 cells; Fig 3G and insets); however, 17/20 cells showed <u>no</u> SkpA recruitment. Together these data are consistent with the idea that Slimb is robustly recruited to the destruction complex via direct or indirect interaction with Axin, and that other E3 ligase components (SkpA and Cul1) are less robustly recruited.

**Figure 3:**
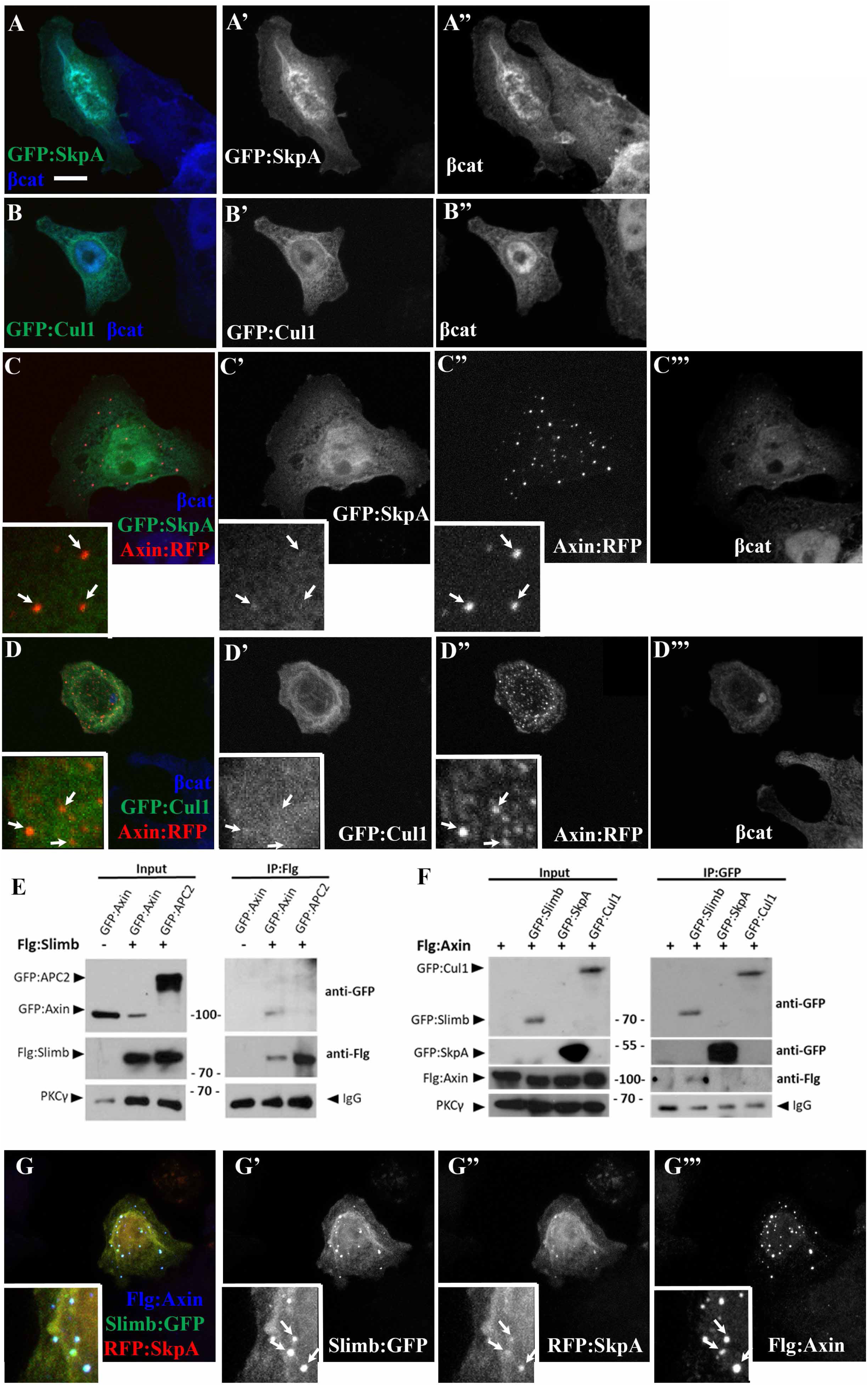
Axin does not recruit Cul1 or SkpA into the destruction complex. (A-D, G) SW480 cells transfected with the indicated *Drosophila* proteins. (A-B) GFP tagged SkpA and Cul1 are both expressed throughout the cytoplasm and nucleus and their expression has little effect on βcat destruction. (C-D) Co-transfection of Axin:RFP with either GFP:SkpA or Cul1 (as marked). Axin is unable to robustly recruit SkpA or Cul1. (E) Testing whether immunoprecipitating Flg-tagged Slimb Co-immunoprecipitated GFP tagged APC2 or Axin. Slimb was able to pull down Axin and not APC2. (F) Testing whether immunoprecipitating GFP-tagged forms of the E3 ligase proteins co-immunoprecipitated Flg tagged Axin. Flg:Axin was only able to coIP with Slimb and not with SkpA or Cul1. All blots are from the same gel, with extraneous lanes removed. (G) Co-expression of Flg:Axin, Slimb:GFP, and RFP:SkpA. Axin is able to robustly recruit Slimb but is less efficient at recruiting SkpA, insets. Arrows point to the same punctum in each channel. Scale bar = 10 μm.

### Axin’s RGS domain is required for efficient Slimb recruitment but its βcat-binding domain is not

Both Axin and Slimb directly bind to βcat, but at different locations on βcat (Xu and Kimelman, 2007). Therefore, some studies suggested that the Axin:Slimb interaction in vivo might not be direct, but instead might be mediated via bridging by βcat (Liu *et al*., 1999), or at least be enhanced by this (Kitagawa *et al*., 1999). APC2 is also able to directly bind to βcat. If Axin solely recruits Slimb via a βcat linker, then APC2 should also be able to recruit Slimb, something not supported by our data (Fig 3C). To further test the hypothesis that Axin recruits Slimb via a βcat bridge, we used an Axin mutant lacking the βcat binding site (AxinΔβcat:RFP; Fig 4A; (Pronobis *et al*., 2017) and co-expressed it with Slimb. If βcat is essential as a bridge between Slimb and Axin, we would expect this Axin mutant to no longer recruit Slimb into Axin puncta. In contrast, if Axin and Slimb can also interact by another means, then Slimb should still be recruited into the puncta. When AxinΔβcat:RFP was expressed alone, it formed cytoplasmic puncta, consistent with the fact that it still contains Axin’s self-polymerization DIX domain (Fig 4C), though it is unable to target βcat for destruction (Pronobis *et al*., 2017). Strikingly, when co-expressed with Slimb:GFP, AxinΔβcat:RFP was still able to robustly recruit Slimb:GFP into puncta (Fig 4F; 19/19 cells showed recruitment), suggesting that the Slimb-Axin interaction is not solely a result of both proteins binding to βcat.

**Figure 4:**
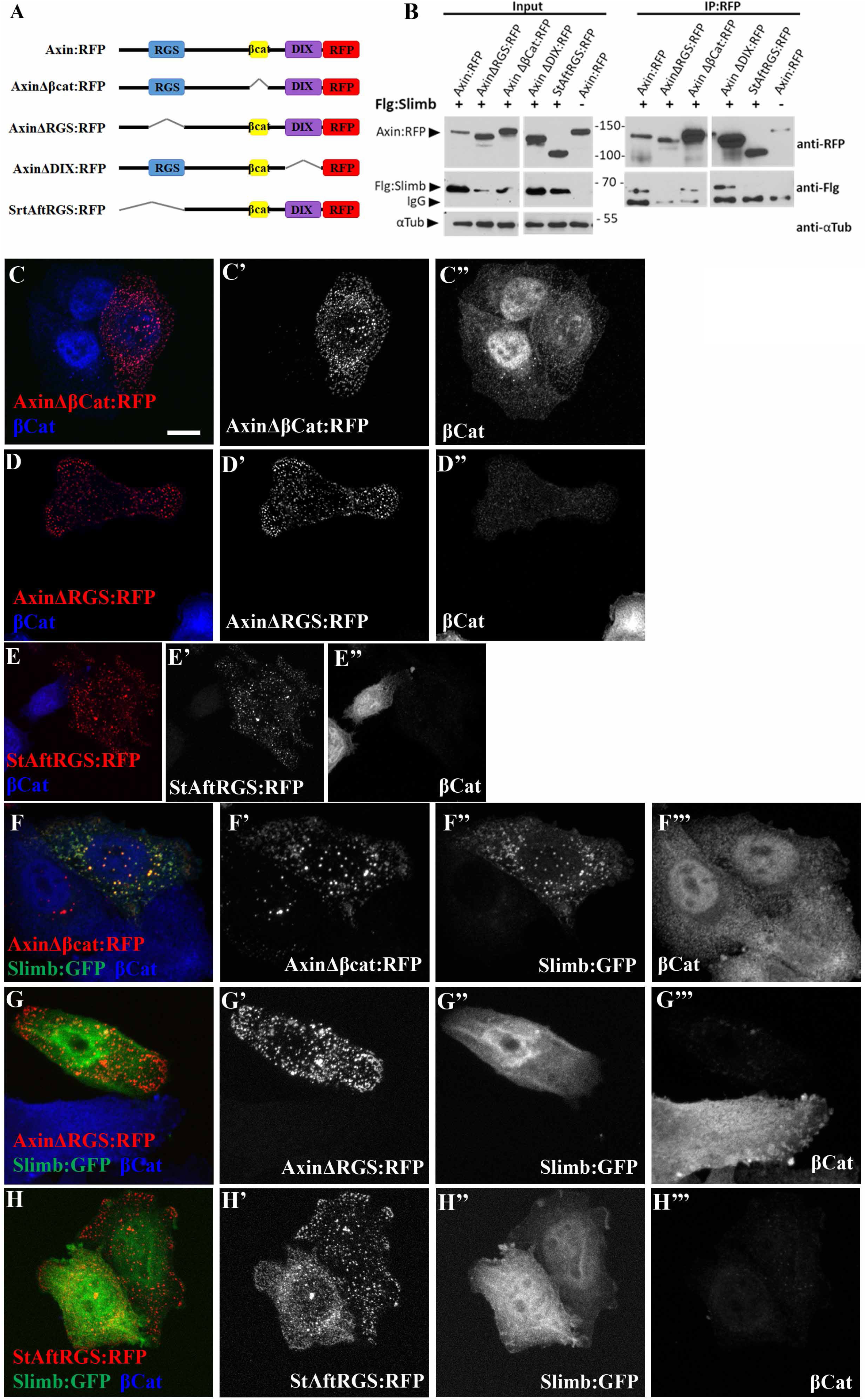
The RGS domain of Axin enhances Slimb recruitment into Axin puncta while the βcat binding site is not essential. (A) Diagram of different Axin mutant constructs used. B) Testing whether Flag-tagged Slimb co-immunoprecipitates with different RFP-tagged Axin mutants. Axin constructs missing the RGS domain are unable to pull down Slimb. All blots are from the same gel, with non-relevant lanes removed. (C-H) Expression in SW480 cells of different Axin mutants, as indicated, alone (C-E) or with Slimb:GFP (F-H). Only AxinΔβcat:RFP is able to robustly recruit Slimb into puncta. Scale bar = 10 μm.

To further investigate which domain(s) of Axin are required for Slimb recruitment, we generated two additional mutants of Axin deleting other domains or regions (Fig 4A): 1) AxinΔRGS:RFP removed the RGS domain, which mediates one of the Axin:APC interactions (Spink *et al*., 2000) and 2) Start After RGS:RFP, which lacks the N-terminal third of Axin. Both mutants retained the ability to form puncta (Fig 4D-E; (Pronobis *et al*., 2017)). Each RFP-tagged Axin mutant was then co-expressed with GFP-tagged Slimb. Both mutants lacking the RGS domain, AxinΔRGS:RFP and Start After RGS:RFP, were diminished in their ability to robustly recruit GFP:Slimb (Fig 4G-H; AxinΔRGS=23/27 cells showed no recruitment; Start After RGS=7/10 cells showed no recruitment). To test this interaction in another way, we IPed Axin mutants tagged with RFP, and assessed if Slimb was co-IPed. Full length Axin, AxinΔβcat:RFP and AxinΔDIX:RFP (Axin lacking its self-polymerization domain; Fig. 4A) all were able to coIP Slimb (Fig. 4B). However, the two Axin mutants lacking the RGS domain did not effectively coIP with Flag:Slimb (Fig 4B). These data suggest that the RGS domain helps mediate the Axin-Slimb interaction.

### Slimb recruitment into the destruction complex can be mediated by either the N-terminal or C-terminal regions of the protein

We similarly asked which part of the multidomain Slimb protein is required for recruitment. We divided Slimb protein roughly in half, separating the N-terminal F-box, which binds SkpA, from the C-terminal WD40 repeats, which dock substrate (Fig 5A). When expressed alone, both halves localized throughout the cytoplasm and nucleus (Fig 5B, D). Strikingly, both halves could be recruited to Axin puncta (Fig 5C,E, arrows in insets), though neither was as robustly recruited (e.g., Fig 5F; the N-terminal half was recruited into Axin puncta in 35/50 cells, the C-terminal half was recruited into Axin puncta in 34/51 cells) as full-length Slim (Fig 1). These data suggest a multipartite binding interaction, consistent with earlier assessment by coIP (Kitagawa *et al*., 1999).

**Figure 5:**
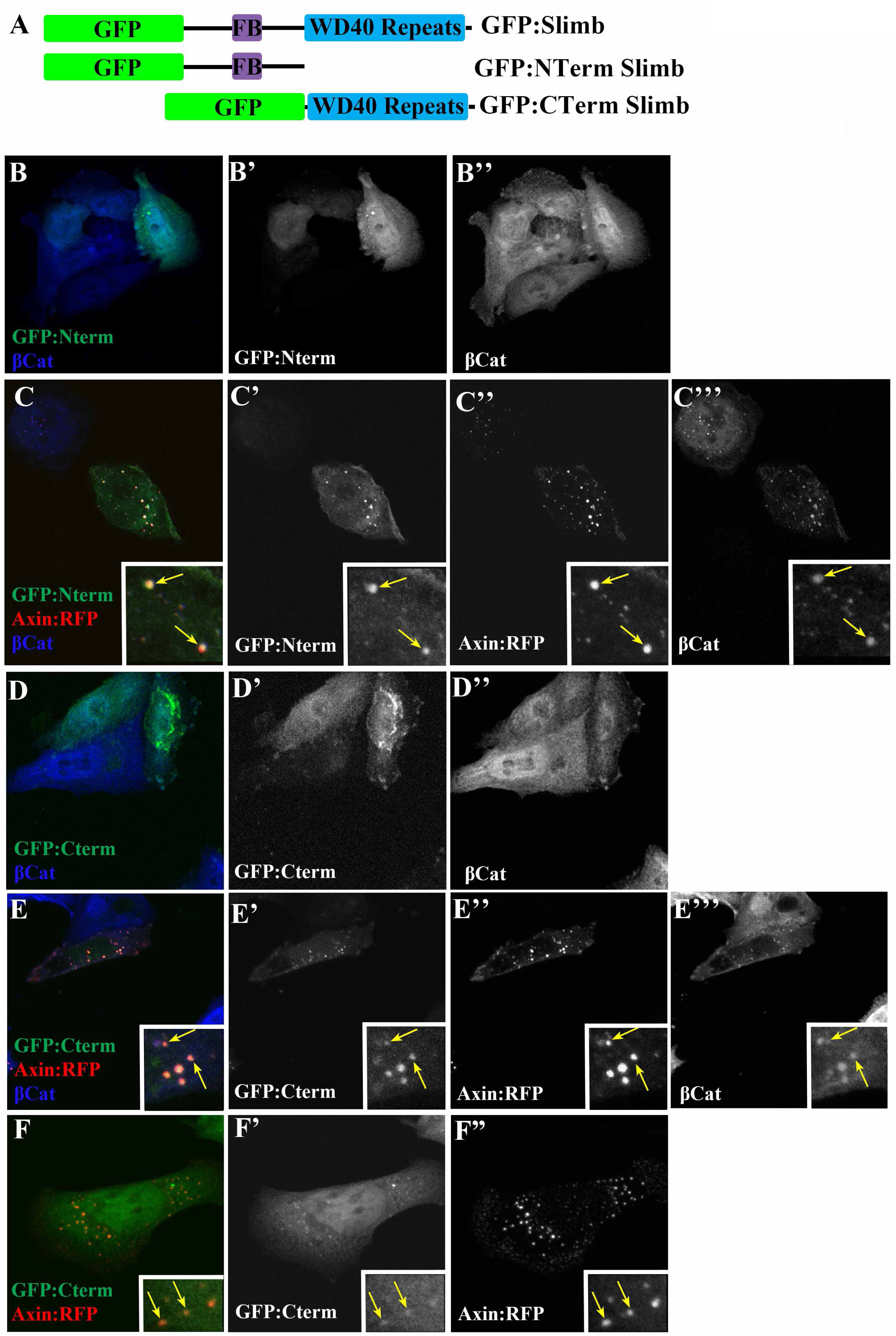
Both halves of Slimb can be recruited by Axin. (A) Diagram of the Slimb pieces created. The N-terminal half contains the F-box domain (FB) with allows binding to SkpA. The C-terminal domain consists mostly of the WD40 repeat domain, which binds to Slimb substrates. (B, D) Both GFP-tagged halves of Slimb are located throughout the cytoplasm and nucleus when expressed alone. (C,E,F) Co-transfection of either half with Axin:RFP reveals that both halves can be recruited to Axin puncta (C,E inset arrows), although the degree of recruitment varies (F).

### Slimb is a dynamic component of the destruction complex

The destruction complex has many of the properties of a biomolecular condensate (Schaefer and Peifer, 2019). One of these is the ability of individual components to rapidly exchange with the cytoplasmic pool— Axin, APC2, and Dsh all can move into and out of puncta (Schwarz-Romond *et al*., 2005; Kunttas-Tatli *et al*., 2014; Pronobis *et al*., 2015). This property can be measured using <u>f</u>luorescence <u>r</u>ecovery <u>a</u>fter <u>p</u>hotobleaching (FRAP), in which fluorescently-tagged protein components of a protein complex are photobleached, and exchange with the unbleached cytoplasmic pool is assessed. FRAP analysis provides an assessment of the mobile fraction (the total amount of protein turnover at the recovery plateau). For example, if there were 100 GFP tagged proteins in a punctum, and we observed 30% recovery of the total GFP fluorescence, this would suggest that on average 70 proteins remained in the complex and 30 new proteins entered. FRAP also provides an assessment of the half-time of recovery (t_1/2_), the amount of time necessary to replace half of the total recovered fluorescence. This measure provides turnover rate. Previous analysis revealed that when expressed <u>alone</u>, Axin:RFP is relatively mobile; however when Axin is co-expressed with APC2, FRAP recovery is less complete and takes significantly longer (Pronobis *et al*., 2015). These data suggested that APC2 stabilizes Axin assembly into puncta. In contrast, Dsh co-expression increases Axin exchange (Schwarz-Romond *et al*., 2007b). To gain an understanding of Slimb dynamics in the destruction complex and the effect of APC2 on its dynamics, we first co-expressed Axin:RFP and GFP:Slimb. Slimb behaved as a dynamic component of the destruction complex (Fig 6A,C). Its recovery plateau was ∼50% and it had a t_1/2_ of ∼100 seconds (Fig 6D). In contrast to Axin, Slimb dynamics were not significantly altered when puncta included both Axin and APC2 (Fig 6B-D), suggesting that stabilization of Axin by APC2 does not stabilize Slimb in the destruction complex. These data suggest that Slimb can form a complex with Axin but is readily able to move out of this complex, consistent with the possibility that Slimb shuttles βcat between the destruction complex and the E3 ligase.

**Figure 6:**
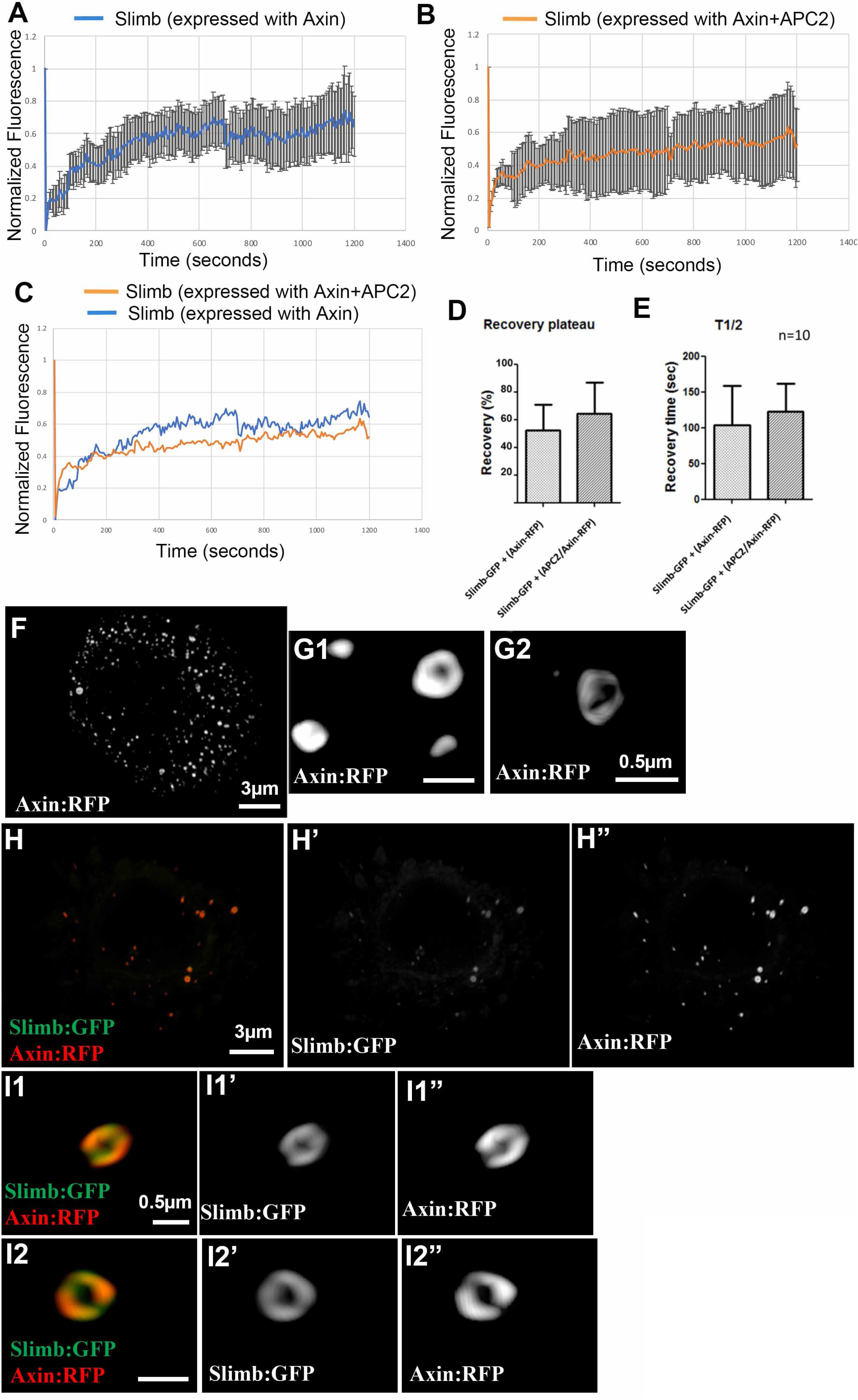
Slimb is a dynamic component of the destruction complex, the turnover of which is not affected by co-expression with APC2 and SIM imaging reveals that Slimb is recruited along Axin cables. (A, B) Slimb:GFP recovery curves when co-expressed with RFP:Axin (A) or with both RFP:Axin and Flag-tagged APC2. Error bars indicate standard errors out of 10 puncta (1 punctum per cell). (C) Recovery curves are similar after co-expression with Axin or with Axin plus APC2. (D) Slimb mobility in puncta is unchanged when expressed with Axin versus APC2 and Axin. F-H. Structured illumination microscopy (SIM) of SW480 cells expressing the indicated constructs, which were directly imaged via the fluorescent tag. (F,H) SIM images of whole cells. Scale bar = 3µm. (G1,G2,I1,I2) Close-up images. scale bar = 0.5 µm. (G1,G2) Axin expressed alone assembles into tight circular cables. (I1,I2) Closeups of puncta in H. Slimb closely localizes along Axin cables.

### Slimb localizes along Axin cables within destruction complex puncta

To further investigate the Axin:Slimb interaction, we visualized this complex by structured illumination microscopy (SIM), which allows increased resolution. When Axin:RFP is expressed alone, puncta contain small polymers of Axin, as we and others previously observed (Fig 6F,G; (Pronobis *et al*., 2015) (Thorvaldsen *et al*., 2015)). In contrast, after co-expression of GFP:APC2 with RFP:Axin, APC2 and Axin form intertwined cables, with an increase in Axin cable size/complexity (Pronobis *et al*., 2015). To explore the relationship of Axin and Slimb within the destruction complex, we co-expressed Axin:RFP and Slimb:GFP (Fig 6H). They colocalized in puncta and the addition of Slimb did not obviously alter average puncta size. Interestingly, Axin and Slimb did not form intertwined cables. Instead Slimb appeared to coat segments of Axin cables (Fig 6I1,I2) These data are consistent with the idea that Axin forms a scaffold upon which Slimb binds. Together, these experiments further our understanding of the mechanisms by which βcat is transferred from the destruction complex to the E3 ligase, thus ensuring its ultimate destruction.

### Exploring how Dsh, Axin and APC cooperate and compete to modulate Wnt signaling in vivo

The data above help illuminate how a functional destruction complex and E3 ligase cooperate to mediate βcat destruction. Wnt signaling can turn down this process, stabilizing βcat and allowing it to enter the nucleus and help activate transcription. We thus next turned to exploring mechanisms underlying this. Cell fate choice in the *Drosophila* embryonic epidermis provides one of the best in vivo models for regulation of Wnt signaling. The GAL4-UAS system likewise offers a superb toolkit to modulate levels of proteins involved in the signaling cascade (Brand and Perrimon, 1993; Duffy, 2002). Using different combinations of GAL4 drivers and UAS-constructs allows titration of protein levels over a wide range. We and others have used this system to probe mechanisms underlying the function of the destruction complex and its downregulation by Wnt signaling.

Co-assembly of Axin and APC is critical to build a functional destruction complex in vivo (e.g. (Mendoza-Topaz *et al*., 2011). Wnt signaling triggers down-regulation of the destruction complex, and the ability to do so depends on relative Axin levels. Thus, while a 3-4 fold increase in Axin levels is tolerated by the developing embryo (Wang *et al*., 2016; Schaefer *et al*., 2018), more substantial increases in Axin levels prevent inactivation of the destruction complex even in cells exposed to the Wnt ligand (Willert *et al*., 1999; Cliffe *et al*., 2003; Schaefer *et al*., 2018). Dsh is a key positive effector of Wnt signaling and its ability to homopolymerize and to heteropolymerize with Axin are critical for downregulating the destruction complex (Schwarz-Romond *et al*., 2007a; Schwarz-Romond *et al*., 2007b; Fiedler *et al*., 2011; Mendoza-Topaz *et al*., 2011). Work in cultured cells suggests Dsh can compete with APC for association with Axin, providing a potential mechanism for Dsh’s role in destruction complex downregulation (Fiedler *et al*., 2011; Mendoza-Topaz *et al*., 2011). However, surprisingly, in *Drosophila* embryos elevating Dsh levels 7-fold has only modest effects on viability and cell fate choices (Cliffe *et al*., 2003; Schaefer *et al*., 2018), suggesting that Dsh may need to be “activated” by Wnt signaling in order to compete for Axin binding.

### Surprisingly, elevating Dsh levels can <u>potentiate</u> the ability of Axin to inhibit Wnt signaling

The simplest versions of the competition hypothesis would predict that elevating Dsh levels would blunt the effects of elevating Axin levels. To test this, we varied the absolute and relative levels of Axin and Dsh. Previous work suggested competition between APC and Dsh for Axin occurs in vivo (Cliffe *et al*., 2003), but those experiments did not assess relative levels of mis-expression, and also did not control for the potential quenching effect on Axin overexpression of driving more than one UAS/Gal4 construct in the same embryo. Our recent work provided mis-expression tools allowing us to control for both these variables, and provide quantitative assessments of relative levels of the two proteins under different conditions (Schaefer *et al*., 2018). We used a strong maternal GAL4 driver line to create robust, uniform, and early maternal and zygotic expression of one transgene (either a GFP-tagged Axin or a Myc-tagged Dsh), and expressed the other UAS-construct zygotically, to allow us to modulate the ratios of Axin and Dsh. We also used a UAS-RFP construct to control for the effects of multiple GAL4 driven transgenes.

We first assessed effects on embryonic viability and cell fate choice. When wildtype embryos secrete cuticle, anterior cells within each segment produce denticles and posterior cells naked cuticle (Fig. 7B, middle). Constitutive activation of Wnt signaling leads to expansion of naked cuticle (Fig 7B, left) while inactivation of Wnt signaling expands denticle cell fates (Fig 7B, right). We used cell fate scoring criteria assessing this which we previously developed (Fig 7C; (Schaefer *et al*., 2018). As we previously found, mild zygotic Axin overexpression (Mat>RFP x Axin; 2-fold increase in Axin levels (Schaefer *et al*., 2018); crosses used and cross nomenclature are in the Methods) had little or no effect on embryonic viability (Fig 7A) or cell fate choices, as assessed by cuticle pattern (Fig. 7C). In contrast, stronger maternal/zygotic Axin overexpression (Mat>Axin x Axin; 9-fold increase in Axin levels; (Schaefer *et al*., 2018) substantially reduced embryonic viability (Fig. 7A), and suppressed Wg-dependent cell fates, thus reducing naked cuticle (Fig 7C; (Schaefer *et al*., 2018). In contrast, increasing levels of Dsh 7-fold (Mat>Dsh x Dsh; (Schaefer *et al*., 2018)) had only modest effects on embryonic viability (Fig. 7A) or cell fate choice (Fig 7C)— the slightly expanded naked cuticle is the cell fate expected for a positive Wnt effector.

**Figure 7:**
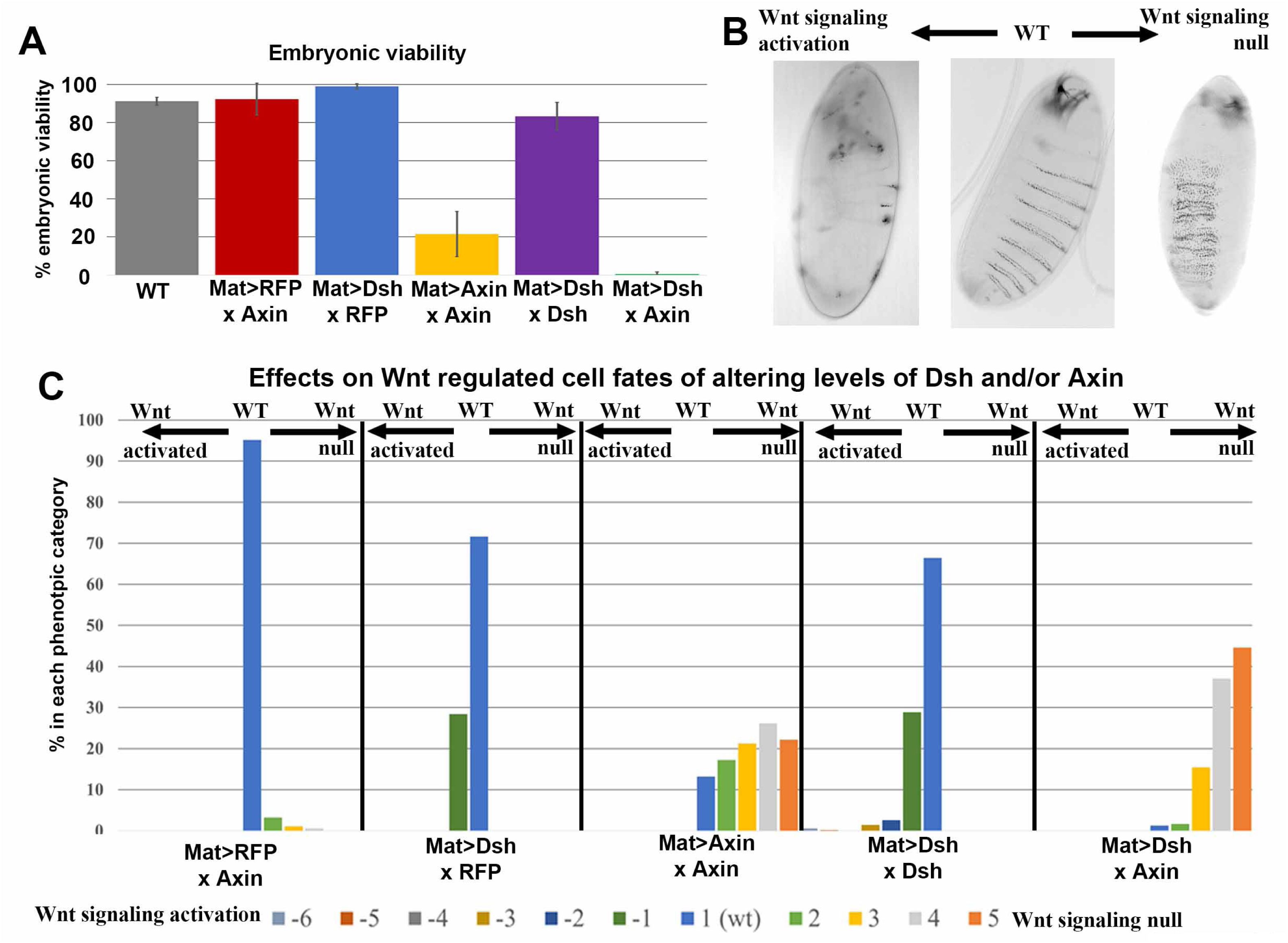
Elevating Dsh levels enhances the ability of Axin to inhibit Wg signaling. (A) Embryonic viability of indicated genotypes. See METHODS for more details on genotype abbreviations. n = 283, 490, 424, 237, 374, 421 respectfully. (B) Representative images of wildtype (center image), with segmentally repeated denticle belts and naked cuticle, flanked by the two most extreme cuticle phenotypes. The left image represents Wnt signaling hyper-activation which results in loss of denticle belts, leaving behind naked cuticle. This is equivalent to phenotype −6 in C. The right image illustrates strong loss of Wnt signaling, inducing loss of naked cuticle and merging of denticle belts. This is equivalent to phenotype 5 in C. (C) Range of cuticle phenotypes observed in dead embryos and hatched larva of genotypes indicated. Categories are as described in (Schaefer *et al*., 2018). Briefly: −6 = Naked cuticle with no denticle belts and large head hole; −5 = naked cuticle with smaller head hole; −4 = small head hole with patches of denticle; −3 = less than ½ of total denticles remain; −2 = more than ½ of denticle are still present, head appears normal; −1 = at most 1 full or parts of 2 denticle belts are absent; 1 = wildtype cuticle phenotype with alternating naked cuticle and denticle belts; 2 = 1-2 merged denticle belts; 3 = 3-4 merged denticle belts; 4 = most denticle belts are merged and mouth parts are still present; and 5 = merged denticle belts with no head.

If Dsh competes with APC for access to Axin, we hypothesized that Dsh overexpression should blunt the effects of Axin over-expression. To make it more likely that Dsh levels would be sufficiently high to effectively compete with Axin, we expressed Dsh maternally and brought Axin in zygotically (Mat>Dsh x Axin). We first examined embryonic viability, and effects on cell fate choice, as assessed by examining cuticle phenotypes. Maternal expression of Dsh alone (Mat>Dsh x RFP) had no effect on embryonic viability (Fig. 7A), and only modest effects on cell fate choice, reflecting occasional mild activation of Wnt signaling (Fig. 7C). However, when we combined maternal expression of Dsh with zygotic expression of Axin (Mat>Dsh x Axin), the result was quite unexpected. This led to essentially complete embryonic lethality (Fig. 7A) and strong <u>suppression</u> of Wnt signaling, as assessed by cuticle pattern (Fig. 7C), the opposite of what we predicted. In fact, the effect on cell fate choice was as strong or stronger than that seen with maternal and zygotic high-level expression of Axin (Mat>Axin x Axin, Fig. 7C). In contrast, the relatively low-level zygotic expression of Axin alone (Mat>RFP x Axin; Fig 7C) did not affect embryonic viability and only occasionally caused mild inhibition of Wnt signaling. This suggested that elevating Dsh levels could potentiate the ability of Axin to inhibit Wnt signaling.

### Elevating Dsh levels enhances the ability of Axin to target Arm for destruction

The direct target of the destruction complex is Armadillo (Arm), the *Drosophila* homolog of βcat. In wildtype embryos, a single row of cells in each segment expresses the Wnt ligand Wg, which moves across the segment, leading to a graded level of signaling (Fig 8A,B). All cells have a pool of Arm at the plasma membrane, bound to E-cadherin to function in βcat’s other role in adherens junctions. However, in cells that do not receive Wg signal, the destruction complex captures most of the remaining Arm and targets it for destruction, thus creating a graded distribution of cytoplasmic/nuclear Arm across the segment, with highest levels centering on the Wg-expressing cells and lowest levels at the most distant cells from the Wg source (Fig 8A,B). To directly assess how altering the relative ratios of Axin and Dsh affect Arm levels, we examined the Arm accumulation in stage 9 embryos, in which Wg signaling is most active. In wildtype embryos, cell rows expressing Wg and Wg-adjacent cell rows have elevated cytoplasmic/nuclear Arm levels (Fig 8B, red arrows), while cells farthest from Wg-expressing cells have low cytoplasmic/nuclear levels of Arm. We quantified these levels by measuring Arm fluorescence intensity in two groups of cells (as in (Schaefer *et al*., 2018): 1-2 cell rows centered on cells expressing Wg (the Wg stripes) and 1-2 cell rows farthest from the Wg-expressing cells (the interstripes). We quantified both relative Arm levels in stripes versus interstripes (Fig 8G) and also the difference in levels between these populations (Fig 8H).

**Figure 8:**
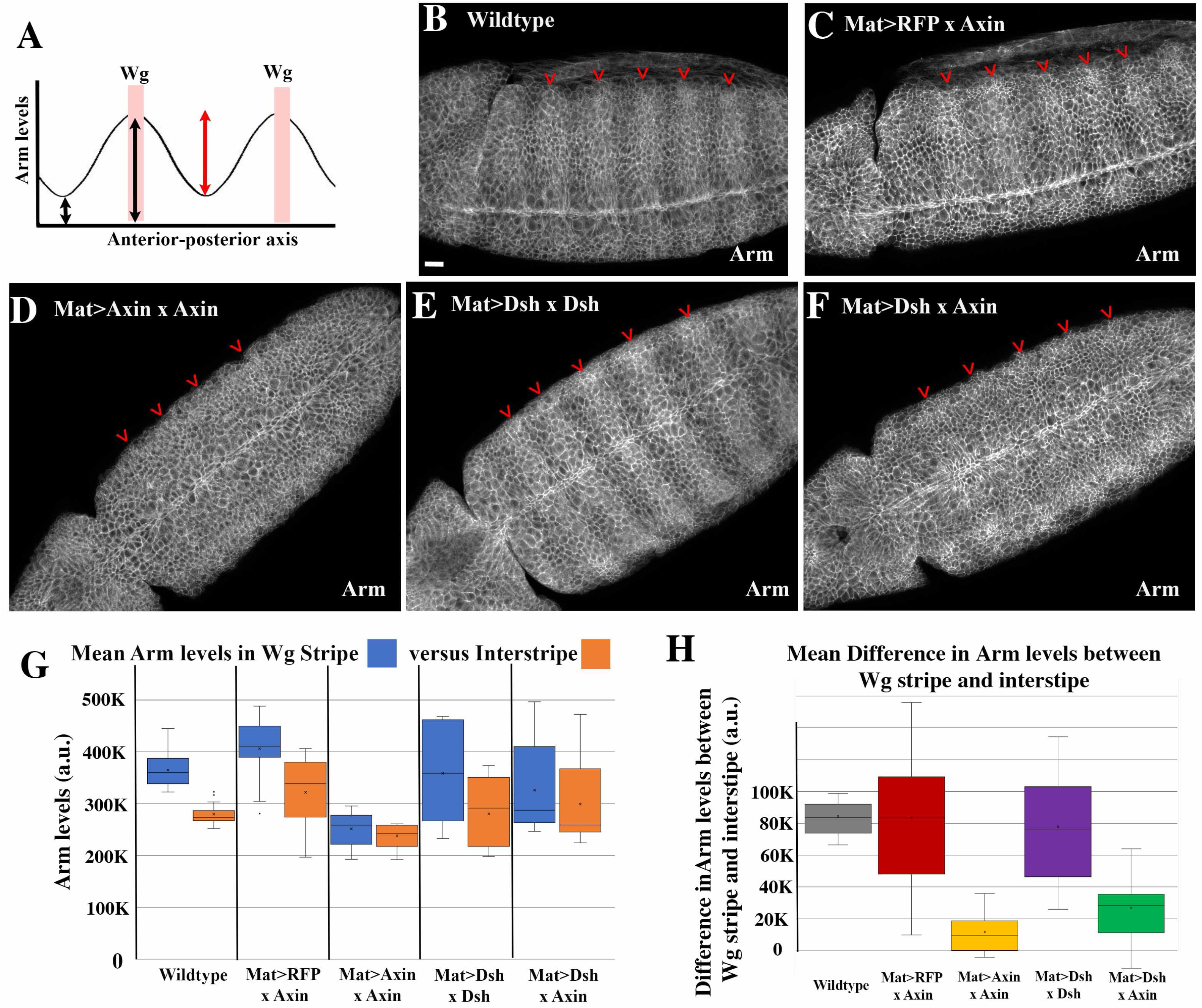
Elevating Dsh levels enhances the ability of Axin to promote Armadillo destruction. (A) Diagram illustrating how Wg-signaling affects Arm accumulation. Two body segments are illustrated. Within each a single row of cells expresses the Wnt ligand Wg, and it forms a graded distribution across the segment, stabilizing cytoplasmic/nuclear Arm in cells that receive it (derived from (Schaefer *et al*., 2018). The closer to the row of cells secreting Wg, the higher the accumulation of Arm in the cytoplasm and nucleus. (B-F) Arm accumulation in Stage 9 embryos, anterior to the left, of the genotypes indicated. Red arrowheads label the row of Wg-expressing cells. (G) Quantification of mean total Arm levels in 2 cell rows with high Wg expression (Wg stripe) versus 2 rows of cells farthest from Wg expressing cells (Interstripe). Box and whisker plots. Boxes cover 25^th^ to 75^th^ percentiles and whiskers the minimum and maximum. Median = middle line, * = average. Error bars = standard deviation of the mean. n = 8 embryos per genotype. (H) Plot of the difference between Arm levels in the Wg stripe vs Interstripe per embryo. Boxes and whiskers as in G’. n = 3 stripes from 8 embryos. (B) In wildtype, Wg stabilizes Arm in a graded fashion with highest levels at the cells that express Wg (quantified in G) and thus a large difference in levels between Arm stripes and interstripes (H). (C) Low level zygotic expression of Axin does not substantially alter Arm stabilization by Wg signaling. (D) High level maternal and zygotic expression of Axin largely abolishes Wg stabilization of Arm. (E) High level maternal and zygotic expression of Dsh does not abolish the graded pattern of Arm accumulation. (F) Combining high level maternal and zygotic expression of Dsh with low level zygotic expression of Axin substantially reduces the ability of Wg to stabilize Arm. Scale bar=15µm.

In wildtype this analysis clearly revealed the stabilization of Arm by Wg signaling in stripe versus interstripe cells (Fig 8B,G,H). In contrast, high level expression of Axin (Mat>Axin x Axin) reduced Arm levels in the Wg stripes (Fig 8D) and thus essentially abolished the difference in levels (Fig 8G,H), as we previously observed (Schaefer *et al*., 2018). In contrast, mild zygotic elevation of Axin levels (Mat>RFP x Axin) had little or no effect on Arm levels (Fig 8C,G,H). However, the same modest elevation of Axin levels had striking effects in embryos that also over-expressed Dsh (Mat>Dsh x Axin)--Arm levels were significantly reduced in cells receiving Wnt signaling (Fig 8F), thus reducing the difference in Arm accumulation between Wnt-ON and Wnt-OFF cells (Fig 8G,H). This resembled the effect of much higher elevation of Axin in embryos <u>not</u> overexpressing Dsh (Fig 8D,G,H; Mat>Axin x Axin), and contrasted with the effects of expressing low levels of Axin alone without elevating Dsh levels (Fig 8C,G,H; Mat>RFP x Axin). As a control, we verified that elevating Dsh levels alone did not significantly disrupt Arm stabilization in Wnt-ON cells or Arm destruction in Wnt-OFF cells (Fig 8E,G,H). Thus elevating Dsh levels enhances the ability of Axin to target Arm for destruction in cells receiving Wnt signals, contrary to our original hypothesis.

As a final assessment of the effect on Wnt signaling of elevating both Dsh and Axin levels, we examined the expression of a Wnt target gene, *engrailed* (*en*). In a wildtype embryo, the two most posterior rows of cells in each segment express Engrailed (Fig 9A,F), and maintenance of this expression requires Wnt signaling (DiNardo *et al*., 1988). To assess effects of our perturbations on Wnt target gene expression, we counted the number of rows of En-expressing cells in stage 9 embryos. Wildtype embryos have two rows of En-expressing cells per segment (Fig 9A,F). In contrast, high level expression of Axin (Mat>Axin x Axin) reduced En expression (Fig 9C,F; average 1 row of En expressing cells per segment; (Schaefer *et al*., 2018). While neither zygotic expression of Axin alone (Mat>RFP x Axin; Fig 9B,F) nor maternal and zygotic expression of Dsh alone (MatDsh x Dsh; Fig 9D,F) had a substantial effect on *en* expression, when we combined maternal expression of Dsh with zygotic expression of Axin (Mat>Dsh x Axin), significantly fewer cells expressed *en* (Fig 9E,F), mimicking the effect high level Axin expression (Mat>Axin x Axin; Fig 9C,F). Thus, whether assessed by embryonic viability, cell fate choice, Arm levels or the expression of a Wnt target gene, elevating Dsh levels can potentiate the ability of Axin to inhibit Wnt signaling.

**Figure 9:**
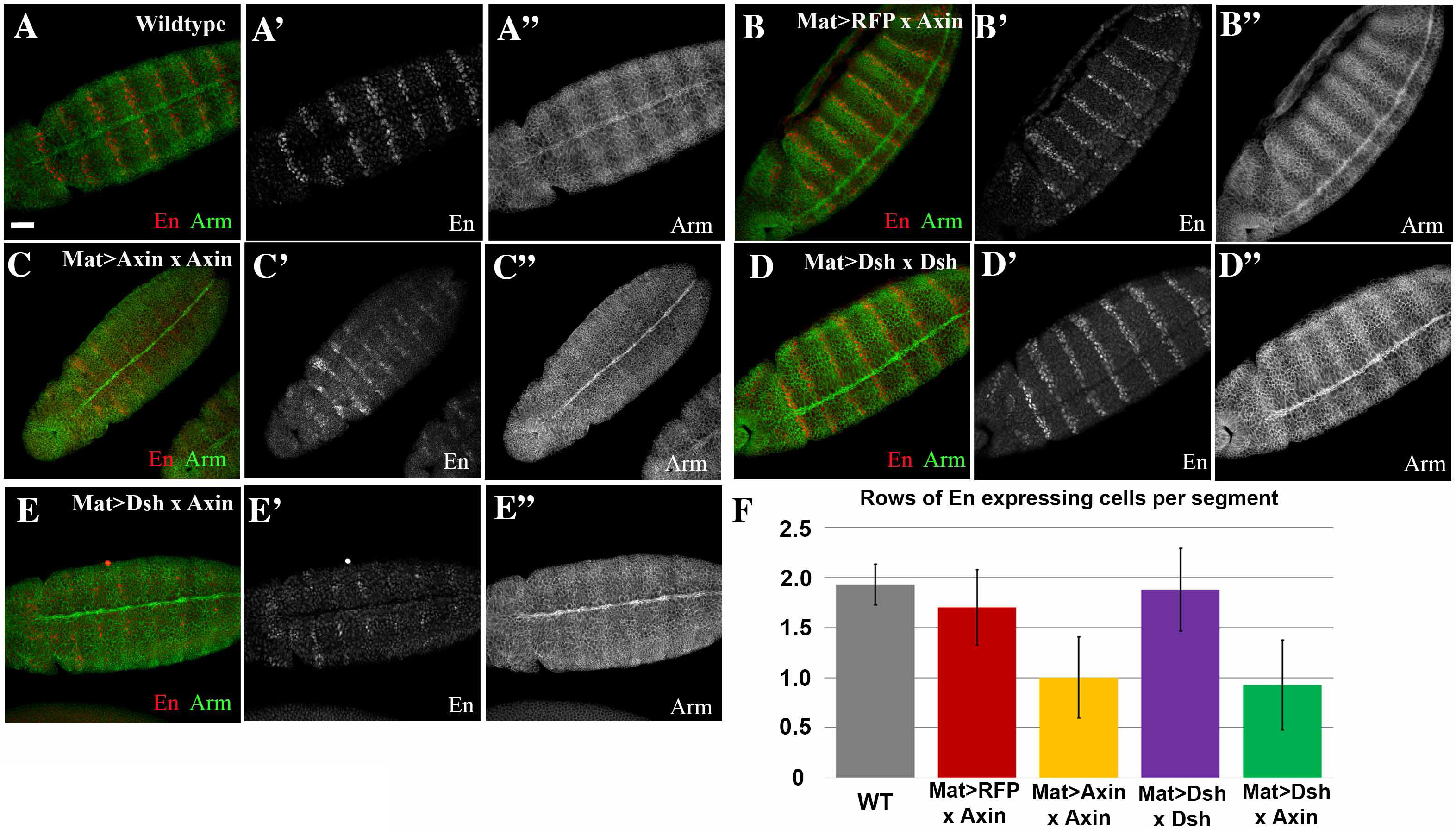
Elevating Dsh levels enhances the ability of Axin to downregulate the Wnt-target gene Engrailed. (A-E) Representative images of Stage 9 embryos for the genotypes indicated, stained for En and Arm. Anterior is to the left. (F) Quantification of the number of En expressing cells per segment. n = 8 embryos for wildtype, and 9 for all other genotypes. Error bars represent the standard deviation of the mean. (A) In wildtype two rows of cells at the posterior of each segment express En. (B) Low level zygotic expression of Axin does not substantially alter the number of cells expressing En (C) High level maternal and zygotic expression of Axin reduces the number of cells expressing En. (D) High level maternal and zygotic expression of Dsh does not substantially alter the number of cells expressing En. (E) Combining high level maternal and zygotic expression of Dsh with low level zygotic expression of Axin substantially reduces the number of En-expressing cells. Scale bar=30µm.

### Elevating Dsh levels in the embryo triggers Dsh assembly into cytoplasmic puncta

In stage 9 *Drosophila* embryos, endogenous Dsh accumulates in the cytoplasm of all cells and is somewhat cortically enriched in cells receiving Wnt signals (Fig 10A, red arrows;(Schaefer *et al*., 2018)-later on, during dorsal closure cortical enrichment increases and Dsh becomes planar polarized to anterior-posterior cell borders in the epidermis (Price *et al*., 2006) Intriguingly, when we previously examined whether endogenous Dsh and Axin:GFP co-localize in puncta, we found overlap in localization in membrane-associated puncta in Wnt-ON cells (Fig. 10B, yellow arrows), but no strong overlap in active destruction complex puncta in Wnt-OFF cells (Fig. 10B, blue arrows; (Schaefer *et al*., 2018). To further explore the effects of Dsh and Dsh/Axin overexpression, we first examined localization of tagged Dsh constructs after over expression. We looked at two different lines in which fluorescent-protein tagged Dsh constructs were driven by the endogenous *dsh* promotor, in a wildtype background. Dsh:GFP2.35 and Dsh:Clover are expressed at levels within a few-fold of endogenous Dsh (Schaefer *et al*., 2018) and Dsh:GFP2.35 is a derivative of a line that rescues the *dsh* null mutant (Axelrod, 2001). In both cases, Dsh accumulated in cytoplasmic puncta in all cells at stage 9 (Fig 10C,D), with some potential reduction in puncta number in Wnt-ON cells (Fig. 10C, red arrows) Similarly, in Mat>Dsh x Dsh embryos, Dsh:Myc accumulates in apical puncta in all cells at stage 9 (Fig 10E), without obvious modulation in levels and localization with respect to cells expressing Wg. Together these data suggest that accumulation in puncta is not solely a property induced by the Myc-tag. The fact that expression of these constructs does not substantially alter Wnt signaling suggests that the puncta do not sequester Axin, consistent with the idea that Dsh needs to be “activated” to interact with Axin. This is particularly intriguing given evidence that Dsh phosphorylation may affect its ability to homo-polymerize (Bernatik *et al*., 2011; Gonzalez-Sancho *et al*., 2013). Intriguingly, the punctate localization of the fluorescent protein-tagged Dsh proteins was stage-specific, as the proteins encoded by Dsh:GFP2.35 and Dsh:Clover accumulated at the cortex in a planar-polarized fashion after dorsal closure (Fig 10F,G), paralleling endogenous Dsh (Price *et al*., 2006). Thus when Dsh levels are elevated, it accumulates in ectopic puncta—it is possible that these puncta also recruit endogenous Dsh, preventing it from assembling with and helping inactivate destruction complexes in Wnt ON-cells.

**Figure 10.**
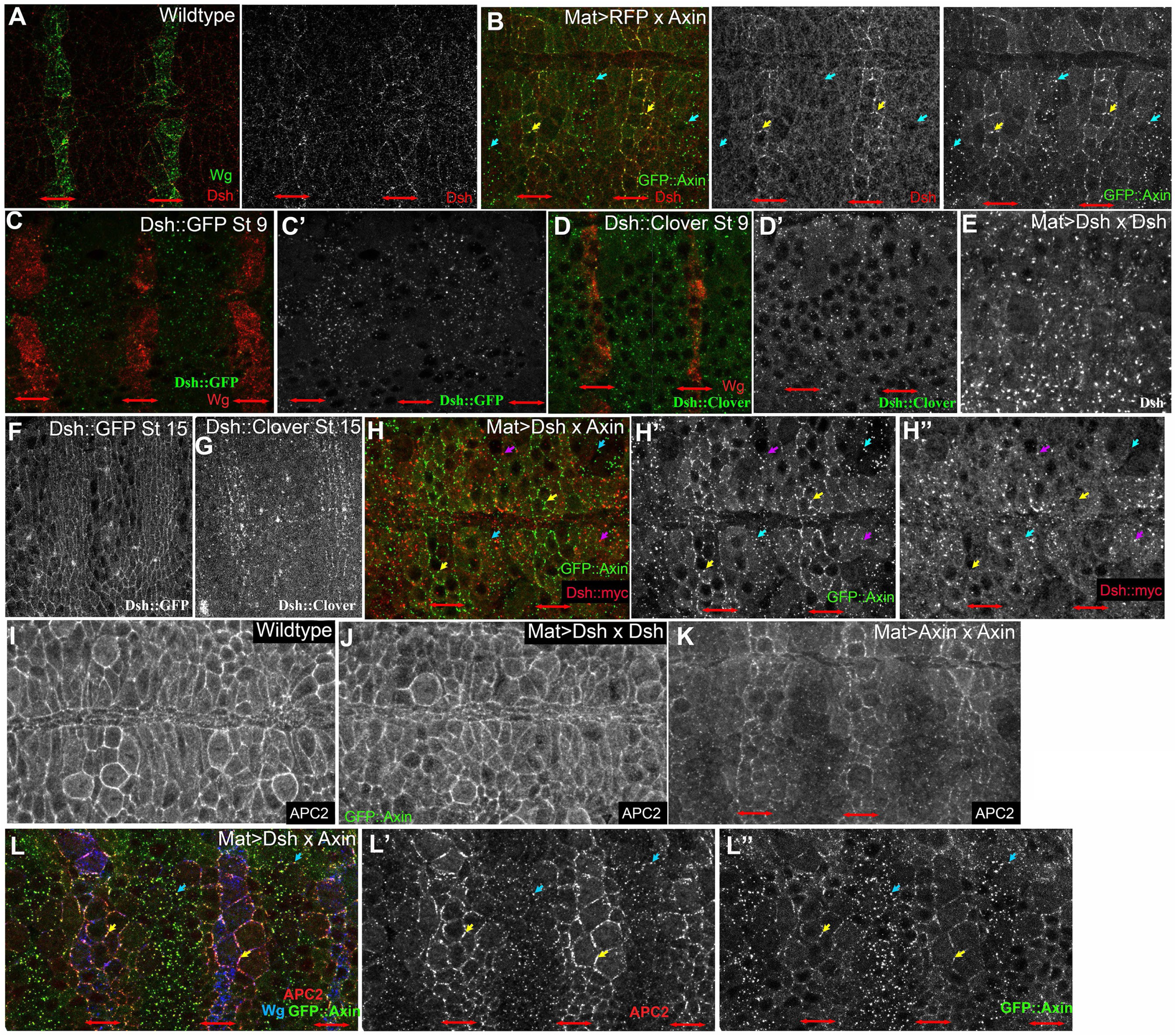
When its expression levels are elevated Dsh forms cytoplasmic puncta but they do not co-localize with Axin. Embryos, anterior to the left, genotypes and antigens indicated. All are stage 9 except F and G, which are stage 15. A. At stage 9 endogenous Dsh is cytoplasmic in all cells with weak membrane enrichment in Wnt-ON cells (in this and subsequent panels WNT-ON cells are indicated by red double-headed arrows). B. Low level overexpression of Axin (Mat>RFP x Axin) enhances recruitment of endogenous Dsh to the membrane of Wnt-ON cells, where it co-localizes with Axin (yellow arrows), but Dsh is not recruited to the cytoplasmic Axin puncta in Wnt-OFF cells (blue arrows). C-D. Two different fluorescent protein tagged Dsh proteins driven by the endogenous promotor both form cytoplasmic puncta at stage 9. E. In Mat>Dsh x Dsh embryos, Dsh:myc forms apical cytoplasmic puncta in all cells. F-G. Two different fluorescent protein tagged Dsh proteins each re-localize to the cortex in a planar-polarized fashion at stage 15, thus resembling endogenous Dsh (Price et al., 2006). H. In Mat>Dsh x Axin embryos (high level Dsh, low Axin), the normal alternating pattern of membrane associated Axin puncta in Wnt-ON cells (yellow arrows) and cytoplasmic Axin puncta in Wnt-OFF cells (blue arrows) is unchanged. Dsh:myc forms cytoplasmic puncta (magenta arrows) but these do not strongly recruit Axin. I. In wildtype embryos APC2 is enriched at the cortex of all cells. J. In Mat>Dsh x Dsh embryos APC2 localization is unchanged. K. Elevating Axin expression (Mat>Axin x Axin) leads to preferential accumulation of APC2 at the cortex of Wnt-ON cells (red arrows). L. In Mat>Dsh x Axin embryos (high level Dsh, low Axin), the recruitment of APC2 to the cortex of Wnt-ON cells is enhanced. Scale bar=15µm.

### Elevating Dsh levels in the embryo does not lead to Axin recruitment into Dsh puncta but does lead to changes in APC localization

We next examined effects of altering Dsh levels on Axin localization. Axin:GFP accumulates in a segmentally varying pattern of localization, with large cytoplasmic puncta in Wnt-OFF cells (Fig 10B blue arrows) and smaller membrane-associated puncta in Wnt-ON cells (Fig 10B, yellow arrows; (Schaefer *et al*., 2018). Membrane relocalization requires Dsh (Cliffe *et al*., 2003). Strikingly, Axin’s segmentally varying pattern was not obviously altered after elevating Dsh levels—Axin:GFP continued to accumulate in large cytoplasmic puncta in Wnt-OFF cells and in smaller membrane-associated puncta and in the cytoplasm of Wnt-ON cells (Fig 10H, blue versus yellow arrows; Mat>Dsh x Axin; compare to Fig 10B). Further, the Dsh puncta that assembled after Dsh:Myc overexpression (Fig 10H, magenta arrows) did not co-localize with the strong Axin puncta in either Wnt-ON or WNT-OFF cells (Fig 10H, yellow and blue arrows arrows), consistent with the idea that “non-activated” Dsh cannot sequester Axin.

We next examined the effect of elevating either Axin or Dsh levels on localization of APC2. Here the effect was more striking. In wildtype embryos, APC2 is cortically enriched in all cells (Fig 10I; (McCartney *et al*., 1999). Elevating levels of Dsh in Mat>Dsh x Dsh embryos does not alter this (Fig 10J). Elevating Axin expression led to a mild increase in cortical enrichment of APC2 in Wnt-ON cells (Fig 10K, red arrows; (Schaefer *et al*., 2018). However, combining elevating Dsh levels with elevating Axin in Mat>Dsh x Axin embryos enhanced the re-localization of APC2 to the plasma membrane of Wnt-ON cells (Fig 10L, yellow arrows), without altering APC localization of Axin puncta in Wnt-OFF cells (Fig 10L blue arrows). Thus elevating Dsh levels also appears to enhance the effects of Axin overexpression on APC2 localization. One speculative possibility is that this could occur because all of the Dsh is assembled into ectopic puncta, preventing it from competing with APC2 for access to Axin.

### SIM imaging suggests relative levels of Axin and Dsh alter their interactions

Our earlier work and that of others illustrated the ability of structured illumination microscopy (SIM) to begin to define the internal structure of the destruction complex, revealing intertwined cables of Axin and APC or of Axin and Tankyrase when these proteins are expressed in cultured human cells (Pronobis *et al*., 2015; Thorvaldsen *et al*., 2015). Like Axin, Dsh forms puncta when expressed in cultured cells. We took a similar approach to better understand the structure of Dsh puncta, utilizing SIM. We tagged Dsh at the N-terminus or the C-terminus with either GFP or RFP, and transfected these constructs into SW480 cells. When expressed alone Dsh forms puncta when tagged with fluorescent proteins either at the N- or C termini (Fig 11A,B)—thus tag localization does not affect the ability of the N-terminal DIX domain to self-polymerize and help drive puncta formation (Schwarz-Romond *et al*., 2005; Schwarz-Romond *et al*., 2007a; Schwarz-Romond *et al*., 2007b; Fiedler *et al*., 2011). Dsh expression had no apparent effect on levels or localization of βcat. Dsh formed two different categories of puncta in different cells: smaller, more spherical puncta (Fig 11A,B) or larger, more complex puncta (Fig 11C,D,G), potentially due to the level of Dsh expression. We then used SIM to look inside these puncta, to see if the Dsh formed an underlying structure within them. When expressed alone, Dsh:RFP in puncta resolved into a loose network of intertwined cables, similar to but more complex than those formed by Axin:RFP expressed alone (Fig 11A,B; (Pronobis *et al*., 2015). When co-expressed, Dsh:GFP and RFP:Dsh largely co-assembled into cables (Fig 11E,E1), suggesting these are not a property of the tag or its location, and consistent with the idea that they co-polymerize. The complexity of the Dsh cables resembled the complexity of the intertwined Axin and APC2 cables formed after co-expression (Fig 12C,D; (Pronobis *et al*., 2015).

**Figure 11:**
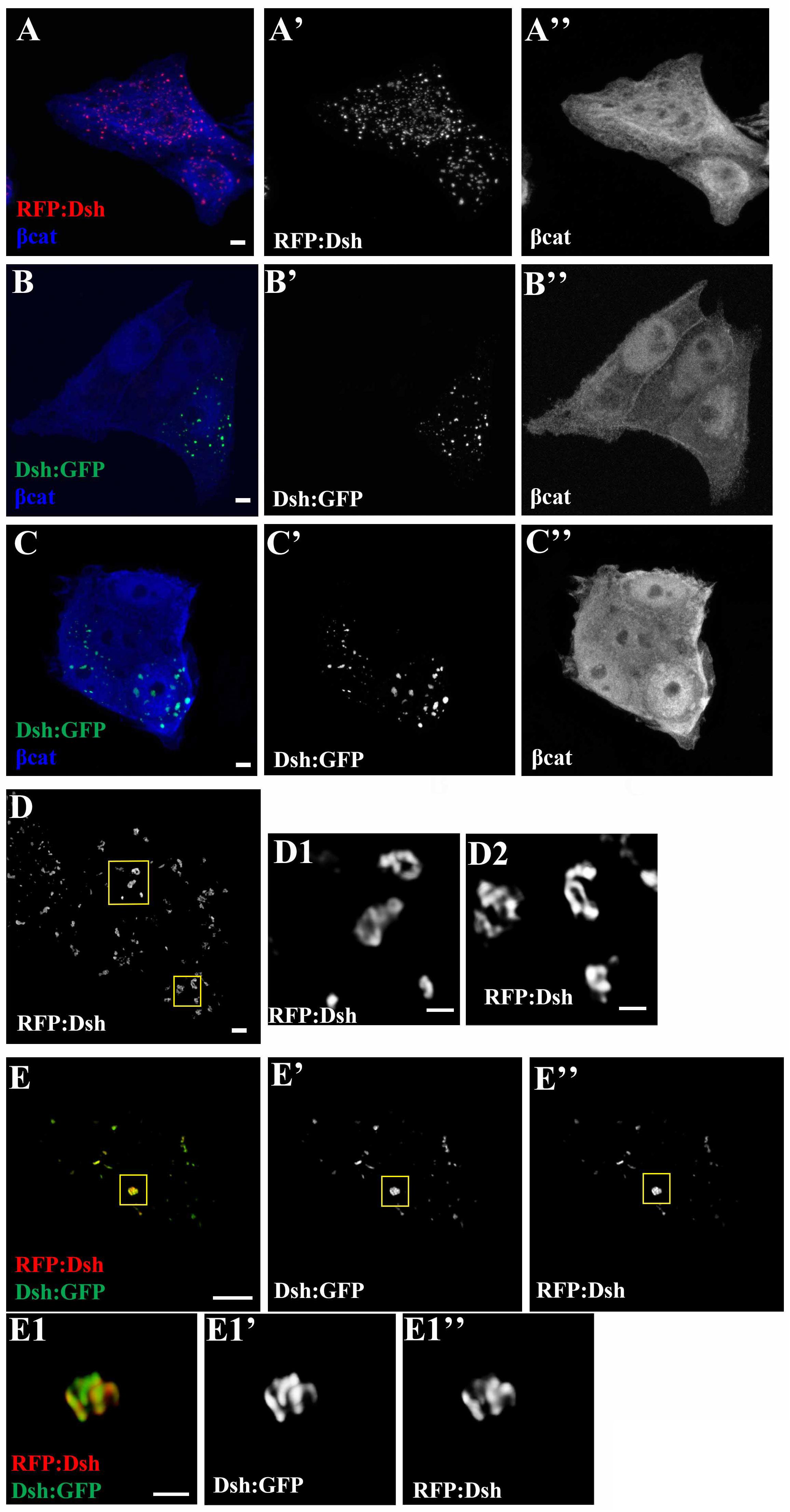
SIM imaging reveals structure inside Dsh puncta. SW480 colorectal cancer cells transfected with the indicated constructs, with fluorescence of the tagged protein imaged directly. (A-C,D,E) Whole cells. Scale bars =10µm (D1,D2,E1) Closeups of puncta. Scale bars=0.5µm (A-B) Confocal images of cells transfected with an N-terminal RFP-tagged Dsh (A) or a C-terminal GFP-tagged Dsh (B), stained for βcat. Both constructs form similar looking puncta, suggesting tag location has little effect on puncta formation. (C) Representative image of two cells, the right of which is expressing higher levels of Dsh. In cells expressing higher levels of Dsh, puncta are larger and less spherical. (D,D1,D2) SIM image of a cell expressing RFP-Dsh, with closeups of several larger puncta. SIM closeups reveal that many puncta have an internal structure of apparent cables. (E,E1) SIM image of a cell co-expressing RFP:Dsh and Dsh:GFP. In closeup of puncta there is intermixing and some co-localization of both versions of tagged Dsh in the internal tangled cables.

**Figure 12:**
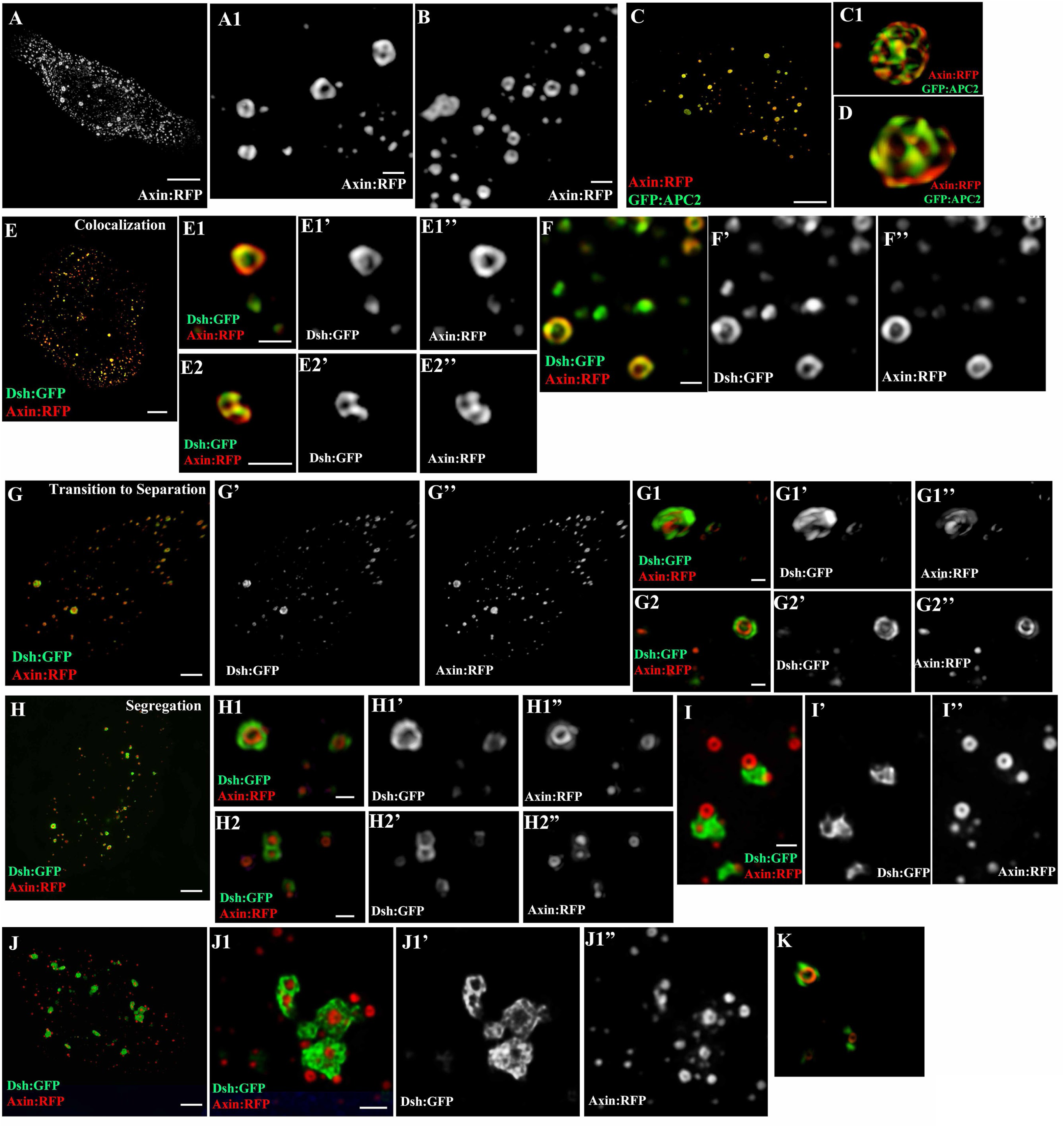
Dsh and Axin can either co-localize in puncta or segregate. SIM imaging of SW480 colorectal cancer cells transfected with the indicated constructs, with fluorescence of the tagged protein imaged directly. (A,C,E,G,H,J) Whole cells. (A1,B,C1,D, E1,E2,F,G1,G2,H1,H2,I, J1) Closeups of puncta. (A-B) When Axin is expressed alone, most Axin puncta contain tight circular cables of Axin (C,C1,D) APC2 co-expression increases the size and complexity of Axin puncta, revealing intertwined cables of Axin and APC2, as we observed in (Pronobis *et al*., 2015). (E-J) Cells expressing both Dsh:GFP and Axin:RFP. (E-F) Representative images from cells with low level Dsh expression. (E1,E2,F) Close-up images of puncta from E or a second similar cell, revealing strong overlap or colocalization of Axin and Dsh. (G-G2) Representative image of a cell in which Axin and Dsh appear to transition into more complete separation. (G1,G2) Close-up of puncta from G. Axin:RFP is more enriched in the center of the punctum, while Dsh:GFP is more enriched in the outer region. (H-J1) SIM image of cells with higher level Dsh expression in which Dsh puncta are larger. (H1,I, J1) Close-ups of punctum from H,J and a third similar cell. Dsh puncta were larger and more complex, while Axin continued for form small circular cables. Rather than co-localizing or strongly overlapping with Axin in puncta, Dsh puncta surrounded (G2,H1,H2,J1) or docked on (I,J1,K) Axin puncta. Scale bars = 3µm for whole cell images and 0.5µm for close-ups

The Bienz group has provided evidence that Dsh and APC compete for access to Axin, and that the Dsh association with Axin inhibits destruction complex function (Cliffe *et al*., 2003; Schwarz-Romond *et al*., 2005; Mendoza-Topaz *et al*., 2011). Both Dsh and Axin can homopolymerize via their DIX domains, and can also heteropolymerize (Schwarz-Romond *et al*., 2005; Schwarz-Romond *et al*., 2007a; Schwarz-Romond *et al*., 2007b; Fiedler *et al*., 2011). Our SW480/SIM system provided an opportunity to explore the Dsh:Axin interaction in vivo at super-resolution. Previous studies revealed that Axin and Dsh can co-localize in cytoplasmic puncta when co-expressed in cells (Fagotto *et al*., 1999; Kishida *et al*., 1999; Julius *et al*., 2000; Fiedler *et al*., 2011). We thus used SIM to examine puncta formed after co-expression of fluorescently-tagged Axin and Dsh. When expressed alone, Axin puncta contain tightly curved Axin filaments (Fig 12A-B; (Pronobis *et al*., 2015). Interestingly when we co-expressed Dsh and Axin, we found that Axin-Dsh puncta interactions varied across a spectrum. In some cells, seemingly those with relatively low levels of Dsh expression, there was strong overlap in localization, with some apparent colocalization (Fig 12E,F). Puncta in this subset of cells formed donut or pretzel shapes (Fig 12E,F), similar to those seen when Axin is expressed alone (Fig 12A,B). However, in cells that appeared to have higher levels of Dsh expression, Axin-Dsh puncta interactions were substantially altered. Axin localization remained largely unchanged, forming primarily small, donut shaped puncta. However, at higher levels of expression, Dsh puncta were larger and more complex (Fig 12G1,H1,I,J), resembling what we saw in cells expressing higher levels of Dsh alone (Fig 11D1,E1). Rather than co-localizing or strongly overlapping with Axin in puncta, we observed complex Dsh structures surrounding (Fig 12G2,H1,H2,J1) or docked on donut-shaped Axin puncta (Fig 12I,J1,K). In many cells, there was little or no co-localization. Together these data suggest that Axin and Dsh can closely interact in puncta, but can also be triggered to segregate from one another by a yet unknown mechanism.

## Discussion

Wnt signaling plays key roles in development and disease, by regulating the stability of its key effector βcat. In the absence of Wnt signals, βcat is phosphorylated by the Wnt-regulatory destruction complex, ubiquitinated by an SCF-class E3 ubiquitin ligase, and destroyed by the proteasome. Binding of Wnt ligands to their Frizzled/LRP receptors stabilizes βcat, via the cytoplasmic effector Dsh. Here we explore two important questions in the field: Is there a direct transfer of βcat from the destruction complex to the E3 ligase, and how does Dsh interaction with the destruction complex protein Axin regulate destruction complex function?

### Defining mechanisms by which βcat is transferred from the destruction complex to the E3 ubiquitin ligase

Regulating the stability of βcat is the key step in Wnt signaling (Peifer *et al*., 1994; van Leeuwen *et al*., 1994). The SCF^Slimb^ E3 ligase was first identified as the relevant E3 regulating βcat levels in 1998 (Jiang and Struhl, 1998; Marikawa and Elinson, 1998). It specifically recognizes βcat after its sequential phosphorylation by CK1 and GSK3 (Hart *et al*., 1999; Kitagawa *et al*., 1999; Liu *et al*., 1999), and the most N-terminal phosphoserine is a key part of the binding site for the F-box protein Slimb/βTrCP (Orford *et al*., 1997; Wu *et al*., 2003). Phosphatase activity in the cytoplasm can rapidly dephosphorylate this residue, raising the question of how βcat is transferred to the E3 ligase without being de-phosphorylated. Earlier work offered two clues. First, βTrCP can coIP with Axin and APC (Hart *et al*., 1999; Kitagawa *et al*., 1999; Liu *et al*., 1999; Li *et al*., 2012), suggesting it may associate, at least transiently, with the destruction complex, providing a potential transfer mechanism. Consistent with this, stabilizing Axin using Tankyrase inhibitors led to co-localization of βTrCP and Tankyrase with the destruction complexes that assemble in response (Thorvaldsen *et al*., 2015). However, it was not clear if this occurred by a direct interaction of βTrCP with destruction complex components, via bridging by phosphorylated βcat, or occurred because other components of the SCF^Slimb^ E3 ligase were recruited more directly, with βTrCP recruited as a secondary consequence. A second clue emerged from analyses revealing that one role for APC is to prevent dephosphorylation of βcat while it is in the destruction complex, protecting the βTrCP binding site (Su *et al*., 2008).

Two plausible models were suggested by this data. In the first, the entire SCF^Slimb^ E3 ligase might be recruited to the destruction complex, allowing direct transfer of phosphorylated βcat between the two complexes. In a second model, βTrCP could serve as a shuttle, binding to phosphorylated βcat at the destruction complex and shuttling it to a place where the E3 could be assembled and ubiquitinate βcat.

We explored interactions of the E3 ligase with the destruction complex using cell biological assays in SW480 cells. We observed ready recruitment of the βTrCP homolog Slimb to destruction complex puncta by Axin, but did not observe recruitment by APC2, consistent with earlier assays by coIP (Kitagawa *et al*., 1999). Slimb recruitment did not require the βcat binding site of Axin, making it less likely that recruitment occurs solely via bridging by βcat. However, it was enhanced by the RGS domain of Axin—future work to assess whether this involves a direct interaction or an indirect one are warranted. There are conserved residues in the RGS domain that are not necessary for the APC-Axin interaction, some of which form a pi helix, and it will be interesting to further explore the function of these residues (Spink *et al*., 2000). Both the region containing the N-terminus plus the F-box of Slimb and that including its WD40 repeats could be separately recruited into Axin puncta, suggesting it may be recruited by multiple interactions—in the case of the WD40 repeats this could include bridging by phosphorylated βcat. Once again, direct binding assays in vitro would provide further insights, building on earlier assays suggesting a multipartite binding interaction (Kitagawa *et al*., 1999). Our super-resolution imaging suggests the interaction between Slimb and Axin is intimate, consistent with direct binding. Our FRAP data, on the other hand, reveal Slimb can come in and out of the complex, similar to the behavior of Axin and APC.

In contrast to the strong recruitment of Slimb to destruction complex puncta, two other core components of the SCF^Slimb^ E3 ligase, Skp1 and Cul1, were not avidly recruited. The occasional recruitment seen could reflect interactions with endogenous βTrCP in the puncta. Co-expression with Slimb slightly enhanced recruitment, but this was still not as robust as the recruitment of Slimb itself. Our IP/Mass spectroscopy data and earlier work from the Mann lab (Hilger and Mann, 2012) are consistent with the presence of all three core SCF^Slimb^ E3 ligase proteins in the destruction complex, but suggest they may be present at lower levels than core destruction complex proteins. One possibility is that Slimb/βTrCP usually acts as a shuttle, but its presence occasionally recruits the other E3 proteins. Another possibility is that the entire SCF^Slimb^ E3 ligase docks on the destruction complex transiently, to accept phosphorylated βcat, ubiquitinate it, and then transfer it to the proteasome. Consistent with this possibility, inhibiting Tankyrase not only stimulates association of βTrCP with Axin, but also leads to recruitment of the proteasome itself to the destruction complex (Thorvaldsen *et al*., 2015)—intriguingly, proteasome inhibition reduces destruction complex assembly, though this effect appears to be indirect, via effects on Axin2 levels (Pedersen *et al*., 2016). Further analyses will be needed to discriminate between these possibilities.

Additional work is also needed to explore how βcat transfer to the E3 ligase is regulated. Direct targeting of βcat to the E3, by fusing the F-box of Slimb to the βcat-binding sites of Tcf4 and E-cadherin, is sufficient to stimulate βcat destruction, independent of the destruction complex (Liu *et al*., 2004), but in vivo the destruction complex plays a critical role. Several pieces of data are consistent with the idea that transfer to the E3 ligase is the step regulated by Wnt-signaling, rather than phosphorylation of βcat, with APC playing an important role (Li *et al*., 2012; Pronobis *et al*., 2015). Further exploration of this process will be welcome.

### Dsh and Axin: a complex interaction

It has been clear for more than two decades that Dsh is a key effector of Wnt signaling (Klingensmith *et al*., 1994; Noordermeer *et al*., 1994). However, its precise mechanisms of action are complex and not fully understood. Current data suggest that Dsh is recruited to activated Frizzled receptors via its DEP domain (Tauriello *et al*., 2012; Gammons *et al*., 2016b). Dsh then helps ensure the Wnt-dependent phosphorylation of LRP5/6 (Bilic *et al*., 2007; Metcalfe *et al*., 2010), leading to receptor clustering, facilitating Axin recruitment, and thus inhibiting GSK3 (Tamai *et al*., 2004; Stamos *et al*., 2014). Dsh homo-polymerization, via its DIX domain, and hetero-polymerization with Axin (Fiedler *et al*., 2011), along with DEP-domain dependent Dsh cross-linking (Gammons *et al*., 2016a), are then thought to lead to downregulation of the destruction complex and thus stabilization of βcat.

Intriguingly, in *Drosophila* embryos Dsh, Axin, and APC are present at levels within a few-fold of one another (Schaefer *et al*., 2018). Many current models suggest that relative ratios of these three proteins are critical to the signaling outcome, with APC and Dsh competing to activate or inhibit Axin, respectively. Consistent with this, substantially elevating Axin levels in vivo, using *Drosophila* embryos as a model, renders the destruction complex immune to downregulation by Wnt-signaling (Willert *et al*., 1999; Cliffe *et al*., 2003). Subsequent work revealed that the precise levels of Axin are critical —elevating Axin levels by 2-4 fold has little effect, while elevation by 9-fold is sufficient to constitutively inactivate Wnt-signaling (Wehrli *et al*., 2000; Wang *et al*., 2016; Schaefer *et al*., 2018). One might then predict that elevating Dsh levels would have the opposite effect, sequestering Axin and thus stabilizing βcat and activating Wnt-signaling. While very high levels of Dsh overexpression can have this effect (Wehrli *et al*., 2000; Cliffe *et al*., 2003), we were surprised to learn that 7-fold elevation of Dsh levels only had a subtle effect on Wnt signaling, consistent with the lack of effect on embryonic viability (Schaefer *et al*., 2018). Our data further suggested that Dsh is only recruited into Axin puncta in cells that received Wg signal, in which puncta are recruited to the plasma membrane, although seemingly similar levels of Dsh were present in Wnt-OFF cells (Schaefer *et al*., 2018). This opened the possibility that a Wnt-stimulated activation event, such as Dsh phosphorylation (e.g. (Yanagawa *et al*., 1995; Gonzalez-Sancho *et al*., 2004), might be required to facilitate Dsh interaction with Axin and thus Axin inactivation. In this scenario, elevating Dsh levels in cells without this activation event –e.g., Wnt-OFF cells--would not alter signaling output.

The simplest versions of the antagonism model, involving competition between formation of Axin/APC vs Axin/Dsh complexes, would also suggest that elevating Dsh levels should alleviate effects of activating Axin. We tested this directly, expressing high levels of Dsh maternally, and lower levels of Axin zygotically. We anticipated that elevating Dsh levels would blunt the effects of elevating levels of Axin. Instead, we got a substantial surprise: elevating levels of Dsh <u>enhanced</u> the ability of Axin to resist turndown by Wnt signaling, thus leading to global activation of the destruction complex and inactivation of Wnt signaling. This was true whether we assessed effects on cell fate choice, Arm levels or expression of a Wnt-target gene. Intriguingly, elevating Dsh levels also appeared to enhance the ability of low level Axin expression to alter Axin:APC interactions in Wnt-ON cells—this might provide a clue to an underlying mechanism.

What could explain this paradoxical result? In our view, the most likely explanation is that Wg-dependent “activation” of Dsh is required for it to interact with and thus downregulate Axin. Consistent with this, Dsh phosphorylation can regulate its ability to homopolymerize (Bernatik *et al*., 2011; Gonzalez-Sancho *et al*., 2013). By elevating Dsh levels, we may have exceeded the capacity of this activation system. High levels of “non-activated” Dsh, while unable to interact with Axin, might still interact with other key proteins involved in destruction complex downregulation, sequestering them in non-productive complexes. For example, Dsh can bind casein kinase I (Peters *et al*., 1999; Sakanaka *et al*., 1999; Kishida *et al*., 2001), which is essential for βcat phosphorylation and destruction. With key proteins sequestered, the system might become less able to inactivate the slightly elevated levels of Axin present, thus leading to constitutive activity of the destruction complex. In this scenario, it is not the relative levels of Axin and Dsh that are key, but the relative levels of Axin and “active Dsh”. Elevating Dsh levels may also lead it to preferentially associate with itself, as we observed in SW480 cells—this could recruit endogenous Dsh away from its normal localization with the destruction complex, thus preventing it from participating in inactivating Axin. Our earlier experiments in which we varied protein levels also revealed a second paradox. While elevating Axin levels rendered the destruction complex resistant to turndown by Wnt signaling, elevating the levels of APC2 increased sensitivity to turndown (Schaefer *et al*., 2018). Together these results reveal that there are still important features of Wnt signaling in vivo yet to be uncovered. Further cell biological and biochemical experiments in vivo, combined with new mathematical models of the suspected competition would be extremely useful.

Our SIM experiments may also provide insights in this regard. The ability of Axin and Dsh to both homo- and hetero-polymerize means free monomers must make a choice. It is likely this is a regulated choice, though the mechanism of regulation remains unclear. Our SW480 experiments, while overly simple, may provide an illustration of how the homo:hetero-polymerization balance can shift. In cells in which both Axin and Dsh were expressed at relatively low levels, puncta contained both proteins, and internal structure was consistent with some level of hetero-polymerization. In contrast, when levels of Dsh were significantly higher, Axin and Dsh tended to segregate into separate, adjoining puncta, suggesting the balance was shifted to homo-polymerization, though the polymers retaining the ability to dock on one another. This segregation could allow Axin to remain in a functional destruction complex while Dsh in separate puncta sequestered other Wnt regulating proteins, potentially explaining over-activation in of the destruction complex in Mat>Dsh x Axin. Defining the mechanisms that determine the relevant affinities of each protein for itself versus for its partner will be informative. Intriguingly, we observed a similar docking rather than co-assembly behavior when we imaged the puncta formed by Axin and those formed by the Arm repeat domain of APC2 (Pronobis *et al*., 2015)—this may be another example where relative affinities of proteins for themselves versus their binding partners differ.

## Methods

### Fly stocks, embryonic lethality, and cuticles

All fly stocks, crosses, and embryo developmental experiments were performed at 25°C. For this study: *y w* was used as wildtype. The following stocks were obtained from the Bloomington Stock Center: Maternal alpha tubulin GAL4 (referred to as MatGAL4; a stock carrying both of the GAL4 lines in 7062 and 7063), UAS-Axin:GFP (7225), UAS-Dsh:Myc (9453), UAS-RFP (30556). Embryonic lethality assays and cuticle preparations were as in Wieschaus and Nüsslein-Volhard (Wieschaus and Nüsslein-Volhard, 1986). Inhibition and/or over-expression of Wg signaling was assessed by analyzing embryonic and first instar larvae cuticles with the scoring criteria found in (Schaefer *et al*., 2018).

**Cross Abbreviations (Female x Male):**

**Mat>RFP x Axin =** UAS-RFP/MatGAL4; +/MatGAL4 x UAS-Axin:GFP
**Mat>Axin x Axin =** +/MatGAL4; UAS-Axin:GFP /MatGAL4 x UAS-Axin:GFP
**Mat>Dsh x Dsh =** +/MatGAL4; UAS-Dsh:Myc/MatGAL4 x UAS-Dsh:Myc
**Mat>Dsh x Axin =** + /MatGAL4; UAS-Dsh:Myc/MatGAL4 x UAS-Axin:GFP
**Mat>Dsh x RFP =** + /MatGAL4; UAS-Dsh:Myc/MatGAL4 x UAS-RFP

### Embryo Immunostaining and antibodies

Flies were allowed to lay eggs on apple juice/agar plates with yeast paste for up to 7 hours. A paintbrush was then used to collect embryos were in 0.1% Triton-X in water and eggs were dechorionated in 50% bleach. After fixation for 20 minutes in 1:1 heptane to 9% formaldehyde, with 8mM EGTA added to preserve GFP expression, embryos were devitellinized in 1:1 heptane to methanol. Embryos were then washed in methanol followed by 0.1% Triton-X in PBS, then incubated in blocking buffer (1:1000 normal goat serum diluted in 0.1% Triton-X in PBS) for 30 min. Primary antibody incubation occurred overnight at 4°C, embryos were washed in 0.1% Triton-X in PBS, and then incubated in secondary antibody at room temperature for 1 hr. Embryos were mounted in Aqua Polymount (Polyscience). Primary antibodies were: Wingless (Wg, Developmental Studies Hybridoma Bank (DSHB):4D4, 1:1000), Arm (DSHB:N27 A1, 1:75), En (DSHB:4D9, 1:50), APC2 (McCartney *et al*., 1999), 1:1000), and Dsh (Shimada *et al*., 2001); 1:4000).

### Assessing effects on Engrailed expression

To determine the transcriptional output of Wg signaling, En expression was analyzed in stage 9 embryos. Engrailed-antibody stained embryos were imaged on a Zeiss LSM 710 or 880 scanning confocal microscope. Images were processed as in (Schaefer *et al*., 2018). Briefly, FIJI (Fiji Is Just ImageJ) was used to generate maximum intensity projections 8μm thick. The En channel was then thresholded to highlight En-expressing cells. The number of En-expressing cells were counted in three different locations per En stripe in thoracic/abdominal stripes 2 through 5. The number of cells per En stripe was then determined by averaging these three values. Embryos were scored blind. Significance was assessed using a one-way ANOVA test.

### Quantitative analysis of Arm levels in Wg stripes versus interstripes

To calculate the absolute levels of Arm accumulation in cells receiving or not receiving Wg signals, stage 9 embryos were collected and stained as previously described. Each genotype was imaged on the same day under the same microscope settings. To calculate the level of Arm accumulation, we choose a boxed region 100 pixels wide x 30 pixels high, spanning the width of the Wg-expressing cells, and measured the mean gray value of Arm using FIJI. Three Wg stripe regions from parasegments 2 to 4 were measured, and the average Arm value minus the background value from a region outside the embryo was defined as the Wg stripe Arm value. In the adjacent interstripe regions we used the same box size to measure and calculate Interstripe Arm values. We also measured the relative difference in Arm accumulation between the Wg Stripes and Interstripes.

### Statistics

To determine the significance between intragroup values (Stripe vs. Interstripe within the same genotype), a Paired t-test was used. An unpaired t-test was used to determine the significance between intergroup values (e.g.: Stripe vs Stripe of different genotypes). For multiple comparisons, an ordinary one-way ANOVA followed by Dunnett’s multiple comparisons test were applied.

### Cell Culture and transfection

For all cell culture experiments the human colorectal-cancer cell line, SW480, was used. Cells were maintained in L-15 media (Corning) supplemented with 10% heat inactivated FBS and 1x Pen/Strep (Gibco) at 37° C with ambient CO_2_ levels. For transfection of *Drosophila* proteins into SW480 cells, Lipofectamine 2000 (Life Technologies) was used for transient transfection following manufacturer’s instructions. All constructs contained the pCMV-backbone and Drosophila genes were inserted using the pCR8/Gateway protocol (Invitrogen) and tagged with GFP, RFP, or Flag as described in (Pronobis *et al*., 2015).

### Cell Immunofluorescence and Microscopy

Cells grown on coverslips were collected for immunofluorescence 24 hours after transfection. Briefly, cells were washed in PBS and then fixed in 4% formaldehyde for 5 minutes. Cells were then permeabilized with 0.1% TritonX-100 in PBS for 5 minutes. After 30 minutes in block buffer (0.01% NGS in PBS), cells were incubated in primary antibody for 1-2 hours, washed with PBS, and then incubated in secondary for 1-2 hours. Cells were mounted on microscope slides in Aqua polymount (Polyscience). Primary antibodies used: anti-βCatenin (BD Transduction, 1:800) and anti-M2-Flag (Sigma, 1:1000). Immunostained cells were imaged on an LSM Pascal microscope (Zeiss), an LSM 710 (Zeiss) or an LSM 880 (Zeiss). All images were processed using FIJI to create max intensity projections, and Photoshop CS6 (Adobe, San Jose, CA) was used to adjust input levels so that the signal spanned the entire output grayscale and to adjust brightness and contrast.

### Super-Resolution Microscopy

Transiently transfected cells were stained and collected as previously mentioned. Cells were mounted in Aqua polymount (Polyscience) and coverslips were sealed with nail-polish to prevent hardening of the mounting media. Cells were then imaged using the Nikon Structured Illumination microscope. Images were first processed using the Nikon software using the default settings. Images were then further processed using the IMARIS software.

### Cell Immunoprecipitation and Western blotting

Cells were collected in lysis buffer (150 mM NaCl, 30 mM Tris pH 7.5, 1 mM EDTA, 1% Triton-X-100, 10% glycerol, 0.5 mM DTT, 0.1 mM PMSF plus proteinase/phosphatase inhibitors (EDTA-free, Thermo Scientific; as in (Li *et al*., 2012)) approximately 24 hours after transfection. Antibody was added and samples were incubated on a nutator overnight at 4°C. The next day ProteinA-Sepharose beads (Sigma) were added and samples were incubated on a nutator for 2h at 4°C. After washing in lysis buffer, immuno-precipitated proteins were removed from the beads with 2xSDS buffer and run on an 8 or 10% SDS PAGE gel. After washing in lysis buffer, immuno-precipitated proteins were removed from the beads with 2xSDS buffer and run on an 8 or 10% SDS PAGE gel and transferred to a nitrocellulose membrane. Westerns were visualized using x-ray Film or the Typhoon Imager. Primary Antibodies: anti-GFP (JL-8; Clontech, 1:1000), anti-Flag (Sigma-Aldrich, 1:2000), anti-γ-tubulin (Sigma-Aldrich, 1:2000). Secondary Antibodies: IRDye800CW anti-Mouse (Licor 1:10,000); HRP-conjugated anti-mouse (Sigma 1:1000).

### Mass spectrometry

HEK293T cells were transfected with pCMV-Flag-APC2 and stable cell lines where established using puromycin resistance. Immunoblotting was used to confirm expression. M2 anti-FLAG antibody (Sigma) was used to pulldown Flag-APC2. Cells were washed 3 times with PBS and then harvested in RIPA buffer. The lysate was pre-cleared with protein A beads for 2 hours at 4°C before the anti-FLAG antibody was added overnight at 4°C. Next day proteins A beads where added to the samples and incubated for 2h at 4C. Washes were conducted with RIPA buffer. Trypsin digestion was performed to elute proteins/peptides.

Trypsinized peptides were separated by reverse-phase liquid chromatography using a nanoACQUITY UPLC system (Waters Corporation). Peptides were trapped on a 2 cm column (Pepmap 100, 3 μm particle size, 100 Å pore size) and separated on a 25 cm EASYspray analytical column (75 μm internal diameter, 2.0 μm C18 particle size, 100 Å pore size). The analytical column was heated to 35°C. A 150 min gradient from 1% buffer B (0.1% formic acid in acetonitrile) to 35% buffer B at flowing at 300 nl/min was used. Mass spectrometry analysis was performed by an Orbitrap Elite mass spectrometer (Thermo Scientific). The ion source was operated at 2.6 kV with the ion transfer tube temperature set to 300 °C. Full MS scans (300–2000 m/z) were acquired by the Orbitrap analyzer at 120,000 resolution, and data-dependent MS2 spectra were acquired in the linear ion trap on the 15 most intense ions using a 2.0 m/z isolation window and collision-induced dissociation (35% normalized collision energy). Precursor ions were selected based on charge states (+2 and +3) and intensity thresholds (above 1e5) from the full scan; dynamic exclusion was set to one repeat during 30 sec and a 60 sec exclusion time. A lock mass of 445.120030 was used.

Raw mass spectrometry data files were searched in MaxQuant (1.6.2.3) using the following parameters: specific tryptic digestion with up to 2 missed cleavages, carbamidomethyl fixed modification, variable protein N-terminal acetylation and methionine oxidation, match between runs, label-free quantification (LFQ) with minimum ratio count of 2, and the UniProtKB/Swiss-Prot human canonical and isoform sequence database (release 02/2017). A 1% false discovery rate was applied to all protein identifications. The mass spectrometry proteomics data have been deposited to the ProteomeXchange Consortium via the PRIDE [1] partner repository with the dataset identifier PXD016314.

### Fluorescence recovery after photobleaching (FRAP)

FRAP assays were conducted as previously described (Pronobis *et al*., 2015). In short, cells expressing the indicated constructs were imaged 24-48h after transfection using an Eclipse TE2000-E microscope (Nikon, Japan). Movies were taken at 1 frame/3 s or 1 frame/6 s for 20 min and bleaching was conducted for 8 s with 100% laser power. Movies were processed using the FRAP analyzer in ImageJ. The bleached area and the cell were outlined, background was subtracted and the movie was processed with FRAP profiler. Values were normalized and recovery plateau and standard error were calculated by averaging 10 movies. For t_1/2_ values were processed in GraphPad (La Jolla CA) using non-linear regression (curve fit)-one phase decay. t_1/2_ values of 10 movies were averaged and standard error calculated.

## Author contributions

K.N. Schaefer, M. Pronobis, and M. Peifer conceived the study. K.N. Schaefer led the experimental team. K.N. Schaefer analyzed the effects of Dsh/Axin co-expression, with assistance from C.E. Williams and analysis of Dsh localization by S. Zhang. K.N. Schaefer and M.I.Pronobis analyzed the interactions of the destruction complex with E3 ligase proteins, with help from L. Bauer, M.I Pronobis carried out mass spectroscopy in collaboration with F. Yang and M.B. Major, and D, Goldfarb analyzed the resulting data, All other experiments were carried out by K.N. Schaefer. K.N. Schaefer, M. Pronobis, and M. Peifer wrote the manuscript with input from the other authors. CRediT Taxonomy is as follows: Conceptualization: K.N. Schaefer, M.I. Pronobis, and M. Peifer. Investigation: K.N. Schaefer, M.I. Pronobis, C.E. Williams, L. Bauer, F. Yang, M.B. Major, D, Goldfarb, and M. Peifer. Supervision: K.N. Schaefer and M. Peifer. Writing-Original Draft Preparation: K.N. Schaefer, M.I. Pronobis, and M. Peifer. Writing: Review and Editing: K.N. Schaefer, M.I. Pronobis, C.E. Williams, L. Bauer, F. Yang, M.B. Major, D, Goldfarb, and M. Peifer. Funding acquisition: K.N. Schaefer and M. Peifer

## Acknowledgements

Thanks to M. Bienz, Y. Ahmed, J. Axelrod, T. Uemura, the Bloomington Drosophila Stock Center and the Developmental Studies Hybridoma Bank for key reagents, E. Thornton-Kolbe for lab management, T. Perdue for help with confocal and superresolution microscopy, and A. Spracklen, lab members for helpful advice and suggestions on the manuscript. This work was supported by NIH R35 GM118096 to M.P. K.N.S. was supported by NSF Graduate Fellowship DGE-1650116.

